# SATB1 maintains naive-like identity in antiviral CD8□ T cells by limiting chromatin remodelling at effector gene loci

**DOI:** 10.1101/2025.07.27.667094

**Authors:** Jason K.C. Lee, Adele Barugahare, Ioanna Gemund, Jasmine Li, Jessie O’Hara, Michael Chopin, Daniel Thiele, Maryam Azme, Thomas Bruer, Nicole L. La Gruta, Mark D. Beyer, Joachim Schultze, Brendan E. Russ, Stephen J. Turner

## Abstract

Optimal CD8^+^ T cell differentiation requires the engagement of transcriptional programs that drive effector phenotypes and function, whilst also shutting down transcriptional programs that maintain the naïve state. While the distinct factors that underpin each state are well studied, the molecular mechanisms that control the switch from the naïve to effector state are not fully understood. Utilising integrated analysis of single-cell genomic data, we identified CD8^+^ T cell effector transcriptional networks that are restrained in the naïve state by the chromatin binding protein Special AT-rich sequence binding protein-1 (SATB1). Utilising a SATB1-Tg model, whereby activated CD8^+^ T cells are unable to down regulate SATB1 upon activation, we observed limited effector differentiation and an inability to engage effector programs in response to both primary and secondary influenza A virus infection. Mechanistically, SATB1 limited chromatin remodelling at key gene loci required for the full engagement of the CD8^+^ T cell effector program. Hence, SATB1 is a master gatekeeper that maintains the naïve T cell state and whose downregulation is necessary to allow transition from a naïve to effector state upon T cell activation. SATB1 modulation may provide new strategies for improving CD8^+^ T cell longevity and self-renewal capacity in different immunotherapeutic treatments.

## Introduction

Upon acute virus infection, antigen-specific CD8^+^ T cells are activated and engage an autonomous program of proliferation and differentiation that results in the acquisition of lineage specific function^1, 2^. Once infection is cleared, the majority of activated effector CD8^+^ T cells die, leaving behind a small population of T cells that persist and establish long-lived memory^1, 3, 4, 5^. The generation of effector and memory CD8^+^ T cells is characterised by the expression of cytolytic molecules such as Granzyme (GZM) A and GZMB^6, 7, 8^, proinflammatory cytokines such as Interferon (IFN)-γ, TNF^9, 10, 11^ and CCL5^12, 13^, and upregulation of cell surface receptors such as PD-1^14, 15^, CX3CR1 and CCR5^16^.

The observable changes in effector CD8^+^ T cell phenotype and function upon differentiation are intimately linked to extended cellular proliferation^9, 17, 18, 19^. The combinatorial expression of effector molecules has been used to infer the extent of cellular differentiation. For example, the acquisition of IFN-γ, and concomitant loss of TNF, is associated with a more differentiated effector type and loss of a memory potential and is only apparent once a cell has undergone multiple divisions^9, 17, 18^. Expression of GZMs requires extended division and is also linked to a more terminal CD8^+^ T cell effector phenotype^18, 19,20^. Mechanistically, the need for extended cellular division to enable acquisition of key CD8^+^ T cell effector function is thought to reflect chromatin remodelling at effector gene loci to a stable and transcriptionally permissive state^9^. The engagement of effector transcriptional programs upon T cell activation is underpinned by expression of a suite of transcription factors such as TBET, BLIMP1, RUNX3, and IRF4^21, 22, 23, 24, 25^. Hence, T cell activation engages transcxriptional networks that serve to promote acquisition of lineage specific function and phenotypes.

Importantly, effector differentiation is not just a matter of upregulation of TFs that promote CD8^+^ T cell effector function. Adaptive CD8^+^ T cell naivety is actively maintained through expression of naïve specific transcriptional and epigenetic programs, with shutdown of these genetic networks required for full effector differentiation^26, 27^. Expression of factors such as FOXO1, LEF-1, TCF-1 and BACH2 in naïve CD8^+^ T cells promote self-renewal capacity and limit the ability of to engage effector transcriptional programs required for terminal effector differentiation^28, 29, 30^. Down regulation of FOXO1, TCF-1 and BACH2 upon T cell activation is a key step in enabling effector differentiation and is mediated by epigenetic repression^31, 32^ ^33, 34^. We recently demonstrated that BACH2 deficiency, in the absence of T cell activation, results in spontaneous higher order chromatin reorganisation within naïve CD8^+^ T cells to a state that reflects an effector state^27^. Together these data suggest maintenance T cell naivety is an active state that needs to be overcome to enable full effector T cell commitment. Given that these processes potentiate CD8^+^ T cell immunity and the formation of immunological memory, understanding the molecular factors that regulate this process is paramount.

To identify novel molecular factors that regulate the transition of CD8^+^ T cells from a naïve to effector differentiation state, we utilized our OT-I adoptive transfer model to generate a combined profile of single cell epigenetic and transcriptional data from naïve, effector and memory OT-I CD8^+^ T cell populations after IAV infection^35^. SATB1 was identified as a master negative regulator of CD8^+^ effector genes that are typically upregulated upon activation. The inability to downregulate SATB1 upon CD8^+^ T cell activation limited acquisition of key markers of effector differentiation after acute infection by limiting the chromatin remodeling required for engagement of the effector transcriptional program early after activation. This also promoted a more memory-like phenotypic state. Interestingly, SATB1 overexpression did not impact CD8^+^ T cell division after activation demonstrating an uncoupling of cellular proliferation from acquisition of the effector state. Hence, SATB1 is a master regulator that controls the transition of naïve CD8^+^ T cells to a differentiated effector state by restraining the shut-down of the naïve T cell program, and full engagement of the effector program.

### Identification of SATB1 as a master regulator of gene regulatory networks in naïve CD8^+^ T cells

Naïve CD8^+^ T cell activation results in dynamic changes in distinct transcriptional programs to support the transition of naïve CD8^+^ T cells to the effector state. This transition is associated with dynamic changes in chromatin accessibility, with licensing or decommissioning of cell state specific gene regulatory elements, known as transcriptional enhancers (TEs), that control target gene transcription via interactions with differentiation state specific transcription factors^12, 36, 37, 38^. To identify novel genetic regulatory networks and the TFs associated with transitions between? naïve, effector and memory CD8^+^ T cell states, we utilised our OT-I CD8^+^ TCR transgenic adoptive transfer model^35, 36^ to generate scRNA-seq and scATAC-seq data from naïve, effector and memory OT-I CD8^+^ T cells after influenza A virus infection (A/HKx31-OVA) (**Fig. 1a**). Uniform manifold approximation and projection (UMAP) analysis for the scRNA-seq or scATAC-seq data was able to distinguish naïve, effector and memory subsets (**Fig. 1b, c**). To validate the clustering analysis, we assessed the enrichment of gene signatures characteristic of naïve (e.g. *Tcf7, Foxo1, Lef1*), effector (*Gzma, Gzmb, Ifng, Cd44, Ccl4, Klrg1)*, and memory (*Il7ra, Tcf7,* Cx3cr1) CD8LJ T cell subsets aligning with their expected differentiation states (**Supplementary Fig. 1a**). To link the cluster annotations between the scRNA-seq and scATAC-seq datasets, we performed label transfer by correlating gene expression profiles with chromatin accessibility patterns. While scRNA-seq data from naïve CD8LJ T cells mapped confidently onto corresponding scATAC-seq clusters, the mapping of effector and memory subsets was less robust (**Fig. 1d**). This suggests that although effector and memory CD8LJ T cells may share similar chromatin landscapes, their transcriptional profiles can diverge significantly, likely reflecting differences in activation status^12, 27, 36^.

**Figure 1.**
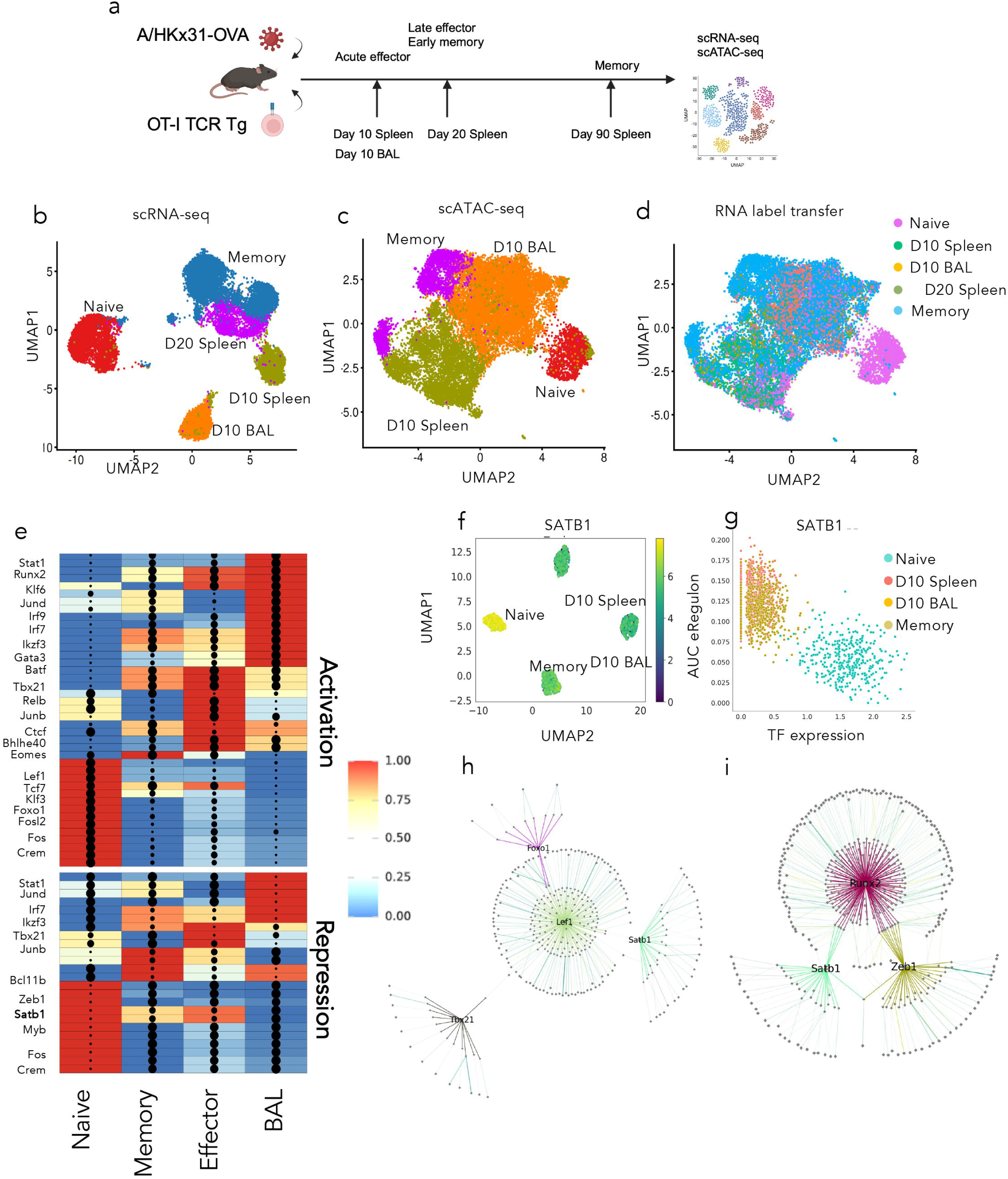
Identification of SATB1 as a transcriptional repressor in naïve CD8^+^ T cells. (**a**) Single cell RNA-sequencing and scATAC-seq was performed on naïve, effector and memory OT-I cells isolated after adoptive transfer into congenic C57BL/6J mice at various timepoints (naïve, effector day 10 (BAL and spleen), day 20; memory, day 90) after infection with A/HKx31-OVA. (**b-d**) scRNA (**b**) and scATAC seq (**c**) data was projected as Uniform Manifold Approximation and Projection with clusters called. (**d**) RNA-seq data was then projected onto the scATAC-seq data using RNA label transfer. (**e**) scRNA-seq and scATAC-seq data was integrated and then analysed using SCENIC+. Heat map/dot-plot showing TF expression and the gene enrichment of eRegulons on a colour scale identified for naïve, effector (BAL/Spleen) and memory OT-I cells. Dots represent the number of genes within a GRN associated with regulatory elements that exhibit enriched TFBS for a given TF. Top of panel are TFs that correlate with transcriptional upregulation of target genes, lower panel are TFs that correlate with transcriptional down regulation of target genes. (**f**) PCA based on GRN enrichment within naïve, effector and memory OT-I cells showing SATB1 expression within each GRN PCA cluster; (**g**) Assessment of SATB1 expression and the correlation with GRNs that exhibit accessible chromatin enriched with SATB1 binding sites **(h**) Visualisation of eGRN networks controlled by LEF1, FOXO1, SATB1 and TBX21 or (**i**) SATB1, ZEB1 and RUNX2

We utilised the SCENIC+ computational tool^39^ to identify transcription factor driven gene regulatory networks (GRNs) enriched in naïve, effector and memory CD8^+^ T cell states (**Fig. 1e, Supplementary Fig. 1b**). This approach integrates differentially open chromatin regions with TF expression, TF binding site (TFBS) enrichment and target gene transcription. TF-associated GRNs can also be correlated with activation (transcriptional activator) or repression of target gene transcription (repressor) (**Fig. 1e**). GRNs identified in naïve OT-I CD8^+^ T cells that were lost upon memory and effector differentiation were positively regulated by TFs, such as TCF1 (encoded by *Tcf7*), FOXO1 and LEF1, all known to positively regulate the naïve CD8^+^ T cell gene program (**Fig. 1e, Supplementary Figure 2**). Similarly, GRNs activated in effector and memory OT-I T cells were driven by TFs known to regulate effector and memory T cell differentiation, and included BATF, TBX21, RUNX3, EOMES and PRDM1 (**Fig. 1e, Supplementary Figure 2**). This approach identified RUNX2, a member of the RUNT transcription family, as a key positive regulator of GRNs uniquely activated in effector/memory T cell states (**Fig. 1e, Supplementary Fig. 3a**) RUNX2 TFBS enrichment has been observed at differentially accessible chromatin regions in effector and memory CD8^+^ T cells^40^ and has been shown to be key for maintenance of virus-specific memory CD8^+^ T cells^41^. Use of RUNX2 reporter mice demonstrated that RUNX2 expression increased in effector and memory IAV-specific influenza A virus cells (**Supplementary Fig. 3b**), and in tumour infiltrating CD8^+^ T cells (**Supplementary Fig. 2c**). Thus, this approach effectively identifies TFs that positively regulate CD8^+^ T cell effector and memory differentiation.

Of particular interest was the identification of TFs that were highly expressed in the naïve state, which acted as transcriptional repressors (**Fig. 1e, bottom panel**). For instance, our data identified that high ZEB1 transcription was associated with transcriptional repression of GRNs in the naïve state (**Fig 1e**). Upon effector differentiation, ZEB1 downregulation correlated with increased transcription of GRNs in effector and memory OT-I (**Fig. 1e, Supplementary Figure 3d, left panel**). This supports an earlier study showing high ZEB1 expression in naïve CD8^+^ T cells, with ZEB1 downregulation linked to CD8^+^ T cell effector differentiation^42^. Interestingly, loss of *Zeb1* transcription from the naïve to effector/memory states inversely correlated with an increase in gene networks regulated by ZEB1, reflecting a role for ZEB1 as a transcriptional repressor (**Supplementary Figure 3d, right panel**). A second factor, SATB1, was also identified as a transcriptional repressor with high expression in the naïve state correlating with decreased GRN transcription (**Fig. 1e, g)**. SATB1 down regulation was observed with effector differentiation (**Fig. 1f, g**) and a concomitant increase in GRN transcription (**Fig. 1e, g**). This indicates that SATB1 may act as a transcriptional repressor serving and acts to maintain the naïve CD8^+^ T cell state^43^. Network analysis demonstrated that SATB1 was linked to GRNs also regulated by LEF1 (**Fig 1i**), a regulator of naïve CD8^+^ T cell identity^30^, but SATB1 was not associated with effector GRNs regulated by TBX21. Interestingly, both SATB1 and ZEB1 were observed to interact with RUNX2 regulated networks suggesting that SATB1 may act to repress RUNX2 regulation of effector differentiation (**Fig 1j**), and could be a potential focus of future studies. Altogether, these data suggest that SATB1 is a regulator of naïve CD8^+^ T cells and acts to limit engagement of effector GRNs.

### SATB1 overexpression limits primary CD8^+^ T cell effector differentiation and acquisition of function

SATB1 dysfunction in naïve CD8^+^ T cells results in a premature effector phenotype prior to activation^43^. We have previously demonstrated that influenza A virus-specific effector and effector memory (T_EM_) CD8^+^ T cells exhibit low levels of SATB1, while naïve and central memory (T_CM_) CD8^+^ T cells expressed high levels of SATB1^43, 44^. Interestingly, TCR activation of naïve CD8^+^ T cells *in vitro* resulted in a transient increase in SATB1 expression at days 2 and 3 after stimulation, with levels starting to decrease by day 4 (**Fig.2a, b**). These data suggest that early upregulation of SATB1 may play a role in limiting effector differentiation early after activation, with subsequent downregulation is required for full transition to an effector state. To examine if over-expression of SATB1 had an impact on cellular proliferation, WT and SATB1-Tg OT-I cells were labelled with cell trace violet (CTV) and activated *in vitro* with WT SIINFEKL (N4) peptide and cell division assessed 3 days later. Importantly, there was no difference in either the extent of cellular division (**Fig 2c, d**), or the extent of activation as measured by CD69 upregulation (**Fig. 2d**). Hence, elevated SATB1 expression early after activation does not impact early proliferative responses.

**Figure 2.**
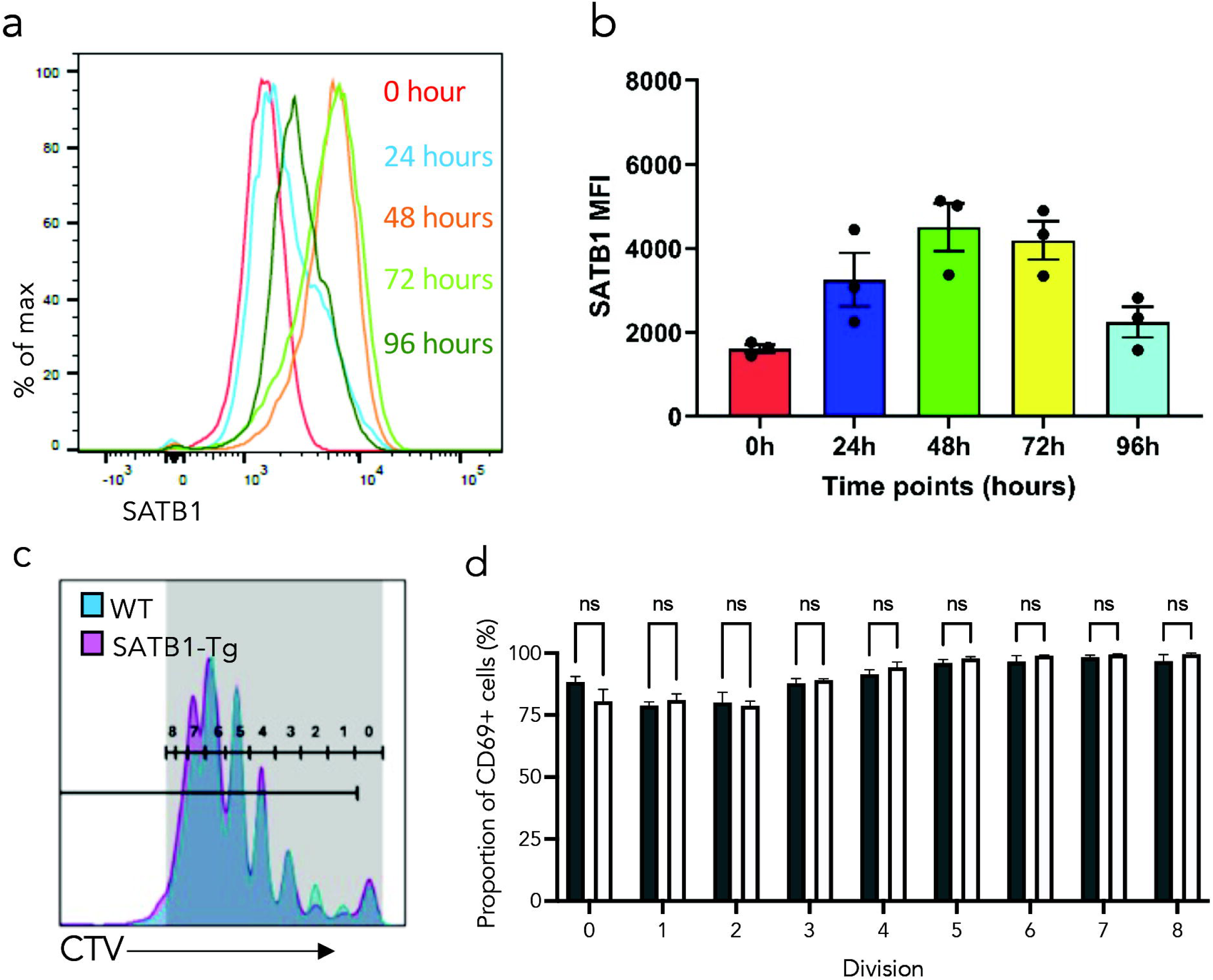
SATB1 is upregulated early after T cell activation without limiting early cell division. (**a, b**) Naïve (CD44^lo^ CD62L^hi^) CD8^+^ T cells were sort purified and stimulated in 96-well plates coated with anti-CD3 (1μg/mL) and anti-CD28 (5μg/mL), for a total of 96 hours in the presence of recombinant human IL-2 (10U/mL). (**a**) SATB1 expression was assessed by flow cytometry at 24-hour intervals after activation and (**b**) SATB1 levels quantitated by measuring mean fluorescence intensity (MFI) (**c**). 2 x 10^6^ naïve (CD44^lo^CD62L^hi^) WT (CD45.1) and SATB1-Tg OT-1 (CD45.1/45.2) CD8^+^ T cells were labelled with Cell Trace Violet dye and equal numbers co-adoptively transferred into recipient C57Bl/6J mice (CD45.2) infected intranasally 3 days prior with 10^4^ plaque forming units of A/HKx31-OVA IAV. (**c**) WT and SATB1-Tg OT-1 were isolated from the mLN 3 days after infection, and the extent of division determined by loss of CTV. (**d**) Dividing cells were co-stained with anti-CD69 to assess the proportion of cells of recently activated CD8+ T cells within dividing WT (grey bars) and SATB1-Tg (white bars) OT-1 CD8^+^ T populations. Bar graphs showing the mean proportion ± SEM of WT OT-1 and SATB1 Tg OT-1 cells expressing CD69 at various divisions.

To examine how over expression of SATB1 impacted CD8+ T cell differentiation during a virus infection, we utilised CD8^+^ T cell-specific SATB1 transgenic mice (SATB1-Tg)^45^ crossed with OT-I TCR Tg mice T cells to examine how the inability to downregulate SATB1 would impact effector CD8^+^ T cell differentiation after IAV infection. WT OT-1 T cells and SATB-Tg OT-I T cells were co-adoptively transferred into recipient congenic mice, followed by i.n. infection with A/HKx31 IAV (**Fig. 2a**). At the peak of the primary effector response, SATB1-Tg OT-I cells exhibited lower prevalence and total numbers compared to WT OT-I cells (**Fig. 2b, c**), particularly in the spleen and mLN. Moreover, SATB1 overexpression skewed the resulting OT-I CD8^+^ effector population towards a less differentiated phenotype with fewer PD-1^hi^ (**Fig. 3d**), and CX3CR1^+^ (**Fig 3e**) cells, with fewer SLECs (CD127^-^KLRG1^+^), and more MPECs (KLRG1^-^CD127^+^) (**Fig. 3f**). SATB1-Tg OT-I cells also expressed much lower levels of TOX, both as a proportion, and on a per cell basis (**Fig. 3g, h**). Importantly, the diminished effector response in SATB1-Tg OT-I CD8+ T cells with a greater proportion of TCF-1^hi^ cells compared to WT controls (**Fig. 3i, j**).

**Figure 3.**
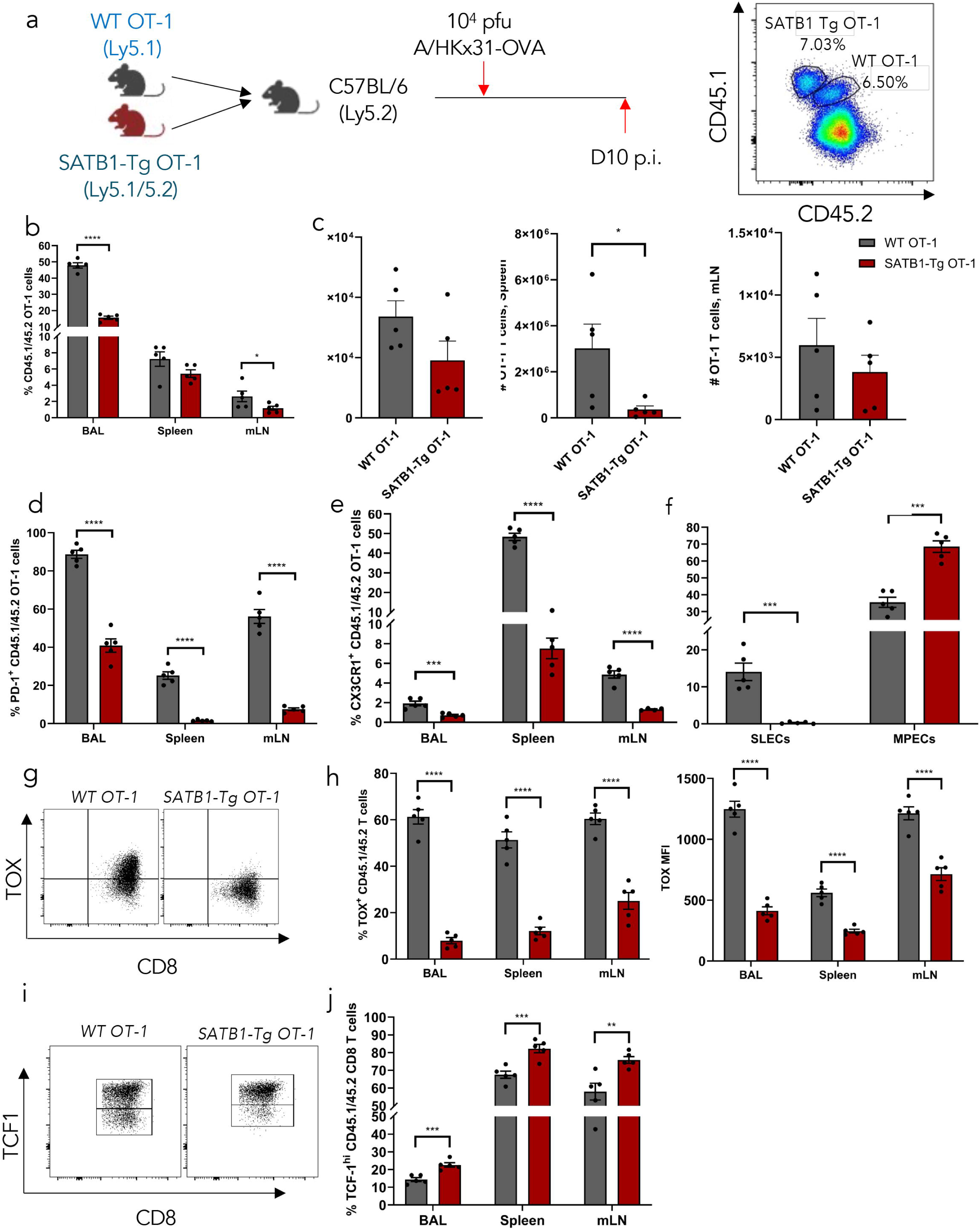
**SATB1 overexpression limits OT-I primary effector CD8^+^ T cell differentiation and expansion**. (**a**) WT (CD45.1/45.2) and SATB1-Tg OT-1 (CD45.1) were co-adoptively transferred into recipient C57Bl/6J mice (CD45.2) followed by intranasal infection with 10^4^ plaque forming units of A/HKx31-OVA influenza virus. Tissues were isolated 10 days after infection and WT OT-1 and SATB1-Tg OT-1 identified based on congenic marker CD45.1/45.2 expression. (**b**) The mean proportion (%) and (**c**) absolute number (#) of WT OT-1 or SATB1-Tg OT-1 in the BAL, spleen and mLN was quantified. The proportion of PD-1^+^ (**d**), CX3CR1^+^ (**e**) and memory precursor (CD127^+^KLRG1^-^) or short-lived effectors (CD127^-^KLRG1^+^ (**f**) OT-1 CD8^+^ T cells were determined for BAL, spleen and mediastinal LN (mLN). The expression of the transcription factors TOX (**g, h**) and TCF-1 (**i-j**) were also determined by flow cytometry. Data is representative of three independent experiments, n=5 with data showing means ± SEM. * *P* ≤0.05, ** *P*≤0.01, *** *P* ≤0.001 and **** *P* ≤0.0001, unpaired student’s *t*-test.

To examine whether SATB1 overexpression also limited endogenous IAV-specific responses, SATB1-Tg mice were infected with IAV and endogenous primary D^b^NP_366_ and D^b^PA_224_ specific CD8^+^ T cell effector responses assessed using tetramers. Endogenous D^b^NP_366_ and D^b^PA_224_-specific CD8^+^ effector T cells demonstrated a more limited IAV-specific CD8^+^ T cell response (**Supplementary Figure 4a-c**). This included fewer KLRG1^+^CD127^-^ SLECs and a greater proportion of KLRG1-CD127^+^ MPECs (**Supplementary Figure 4c**), as well as lower expression of more terminal effector markers such as PD-1, CX3CR1, TOX, and GMZA (**Supplementary Figure 4d-i**). Altogether, these data demonstrate that the inability to downregulate intrinsic SATB1 upon CD8^+^ T cell activation limits both effector T cell expansion and the development of a more differentiated effector phenotype during primary IAV responses.

### SATB1 overexpression decouples secondary CD8^+^ T cell effector expansion from effector differentiation

Given that SATB1 overexpression limited primary effector CD8^+^ T cell responses, we then examined whether this also extended to the establishment of T cell memory and subsequent recall responses. WT and SATB-Tg OT-I cells were co-adoptively transferred into recipient congenic mice, followed by i.n. infection with A/HKx31-OVA and memory CD8^+^ T cell responses assessed at day 60 after infection (**Fig. 4a**). There was no difference in the proportion or number of lung or splenic WT or SATB1-Tg OT-I memory CD8^+^ T cells (**Fig. 4b, c**). Despite no difference in numbers, SATB1-Tg OT-I cells did have a higher proportion of central (T_CM_) vs effector (T_EM_) memory CD8^+^ T cells compared to WT OT-I cells (**Fig. 4d**), again suggesting that SATB1 acts to limit effector differentiation, and consistent with T_CM_ expressing more SATB1^43, 44^. Similar observations were made for endogenous D^b^NP_366_ and D^b^PA_224_ CD8^+^ T cell responses with no significant difference in memory T cell numbers between WT and SATB1-Tg mice. However, SATB1-Tg memory D^b^NP_366_ CD8^+^ T cells were biased towards a T_CM_ vs T_EM_ phenotype (**Supplementary Figure 4j-l**).

**Figure 4.**
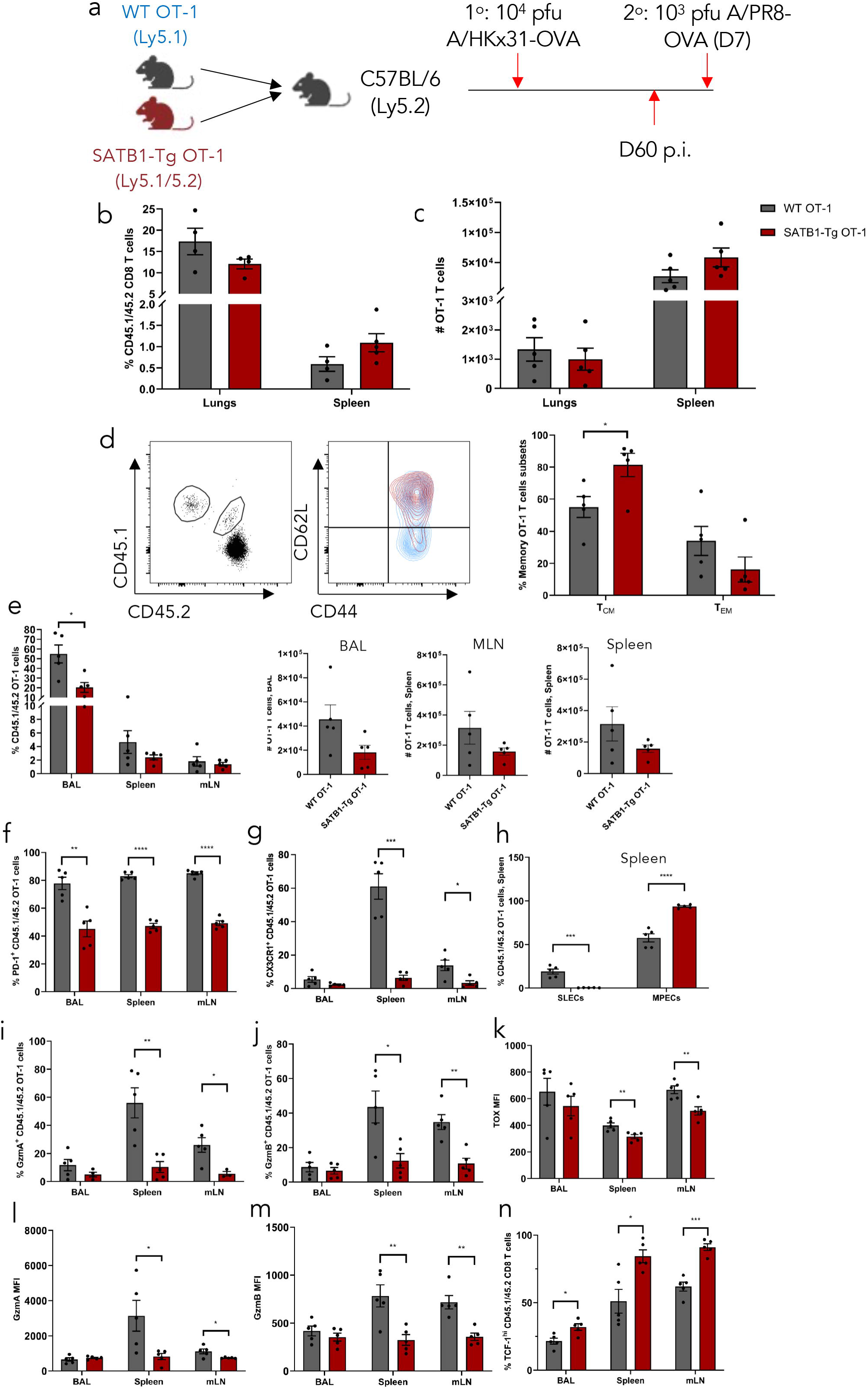
SATB1 overexpression limits phenotypic differentiation but not response magnitude of secondary effector CD8^+^ T cell responses. (**a**) WT (CD45.1/45.2) and SATB1-Tg OT-1 (CD45.1) were co-adoptively transferred into recipient C57Bl/6J mice (CD45.2) followed by intranasal infection with 10^4^ plaque forming units of A/HKx31-OVA influenza virus. A cohort of mice were rechallenged with A/PR8-OVA 60 days after primary infection. Tissues were isolated on day 60 after primary infection (memory) or 7 days after secondary infection and WT OT-1 and SATB1-Tg OT-1 identified based on congenic marker CD45.1/45.2 expression. (**b**) The proportion and (**c**) number of lung and splenic memory WT OT-1 (black bars) or SATB1-Tg OT-1 (white bars) CD8^+^ T cells. (**d**) The proportion of splenic T_CM_ (CD44^hi^CD62L^hi^) vs T_EM_ (CD44^hi^CD62L^lo^) was determined for WT OT-I (CD45.1^+^, blue contour plot, black bars) and SATB1-Tg OT-1 (CD45.1/45.2^+^, red contour plot, white bars). (**e**) Proportion and magnitude of secondary WT OT-I (black bars) and SATB1-Tg OT-1 (white bars from the BAL, spleen and mLN). (**f-h**) Mean proportions of WT OT-I (black bars) and SATB1-Tg OT-1 (white bars) CD8^+^ secondary effectors expressing PD-1 (**f**) CX3CR1 (**g**) and SLEC or MPEC markers (**h**) as described above. Mean proportions (**i-k**) and mean fluorescence intensity (MFI) of WT OT-I (black bars) and SATB1-Tg OT-1 (white bars) CD8^+^ secondary effectors expressing granzyme A (**i, l**), granzyme B (**j, m**) or TOX (**k, n**). Data is representative of 2 independent experiments with n =5 female and represented as mean ± SEM. * indicates *P* ≤0.05, ** *P* ≤0.01, *** *P* ≤0.001 and **** *P* ≤0.0001, unpaired student’s *t*-test.

Secondary infection resulted in largely equivalent recall responses for both WT and SATB1-Tg OT-I cells in the spleen and draining mLN, although fewer effector were evident in the lung airways (**Fig. 3e**). Interestingly, despite no large difference in the proliferative response of memory CD8^+^ T cells, secondary effector SATB1-Tg OT-I CD8^+^ T cells did exhibit a less differentiated phenotype, similar to the primary effector response, with decreased expression of PD-1, CX3CR1, TOX, a lower proportion of SLECs, higher TCF-1 expression and lower GZMA and GZMB expression (**Fig. 3f-n**). Importantly, a similar pattern of diminished secondary effector differentiation was also observed for endogenous SATB1-Tg IAV-specific CD8^+^ T cells with profound decreased expression of PD-1, CX3CR1 and GZMs by SATB1-Tg D^b^NP_366_-and D^b^PA_224_ specific CD8^+^ T cell, despite little or no difference in the magnitude of the overall secondary T cell responses (**Supplementary Fig. 5**). It was of particular interest that sustained SATB1 expression appeared to enable extensive proliferation (**Fig. 2c, d**) but limited acquisition of an effector phenotype, essentially decoupling these attributes.

### SATB1 limits chromatin accessibility at gene loci linked to effector differentiation

We have previously demonstrated that expression of a mutant version of SATB1 that abrogates DNA binding resulted in premature expression of CD8^+^ T cell effector markers within naïve CD8^+^ T cells^43^. This suggests that SATB1 is actively engaged in limiting the acquisition of effector gene signatures. A comparison of transcriptional profiles between WT and SATB1-Tg CD8+ T cells demonstrated while there was little difference in gene signatures in the naïve state, over-expression of SATB1 resulted in diminished effector gene expression in cells isolated 10 days after primary infection. This included effector genes such as *Gzma, Zeb2, Cx3cr1, Ccl5, Pdcd1*, *Runx2, Bach2, Ezh2,* and *Tox* (**Supplementary Fig. 6**). Interestingly, effector SATB1-Tg OT-I cells had higher expression of *Tcf7* and *Lef1*, further supporting a role for SATB1 in limiting effector differentiation. We then performed Assay for Transposase-Accessible Chromatin with high throughput sequencing (ATAC-Seq) to assess the differences in chromatin accessibility between naïve and effector WT and SATB1-Tg OT-I cells (**Fig. 5)**. Clustering analysis highlighted that while little difference was observed in the naïve state, SATB1 overexpression restrained full maturation of chromatin remodelling within effector OT-I CD8^+^ T cells (**Fig. 5a**). This effect was particularly noticeable in terminal effector OT-I cells isolated from the BAL. Analysis of differentiational accessible regions (DARs) between effector WT and SATB1-Tg OT-I cells isolated from the BAL or spleen again demonstrated that SATB1 overexpression largely limited chromatin accessibility, particularly at key effector gene loci (**Fig. 5b**), suggesting that SATB1 maintains a repressive chromatin landscape to limit the effector program. While only 237 DARs separated naïve WT and SATB1 Tg OT-I cells (**data not shown**), there were 1,518 DARs and 1,822 DARs identified between WT and SATB1-Tg effector OT-1 CD8^+^ T cells isolated from BAL and spleen, respectively (**Fig. 5b**). In particular, SATB1 overexpression limited accessibility at a transcriptional enhancer within the *Pdcd1* locus known to be key for maintaining PD-1 expression^46^, and key locations within the *Tox* locus, a key driver of T cell exhaustion^47, 48^ (**Fig 5c, d**). Moreover, SATB1 overexpression also limited accessibility at regulatory regions within Zeb2 and *Ezh2*, factors also known to drive effector CD8^+^ T cell differentiation^31, 42^. Overall, these data suggest that SATB1 maintains a more naïve state by limiting access to key regulatory regions within genes that promote terminal effector differentiation.

**Figure 5.**
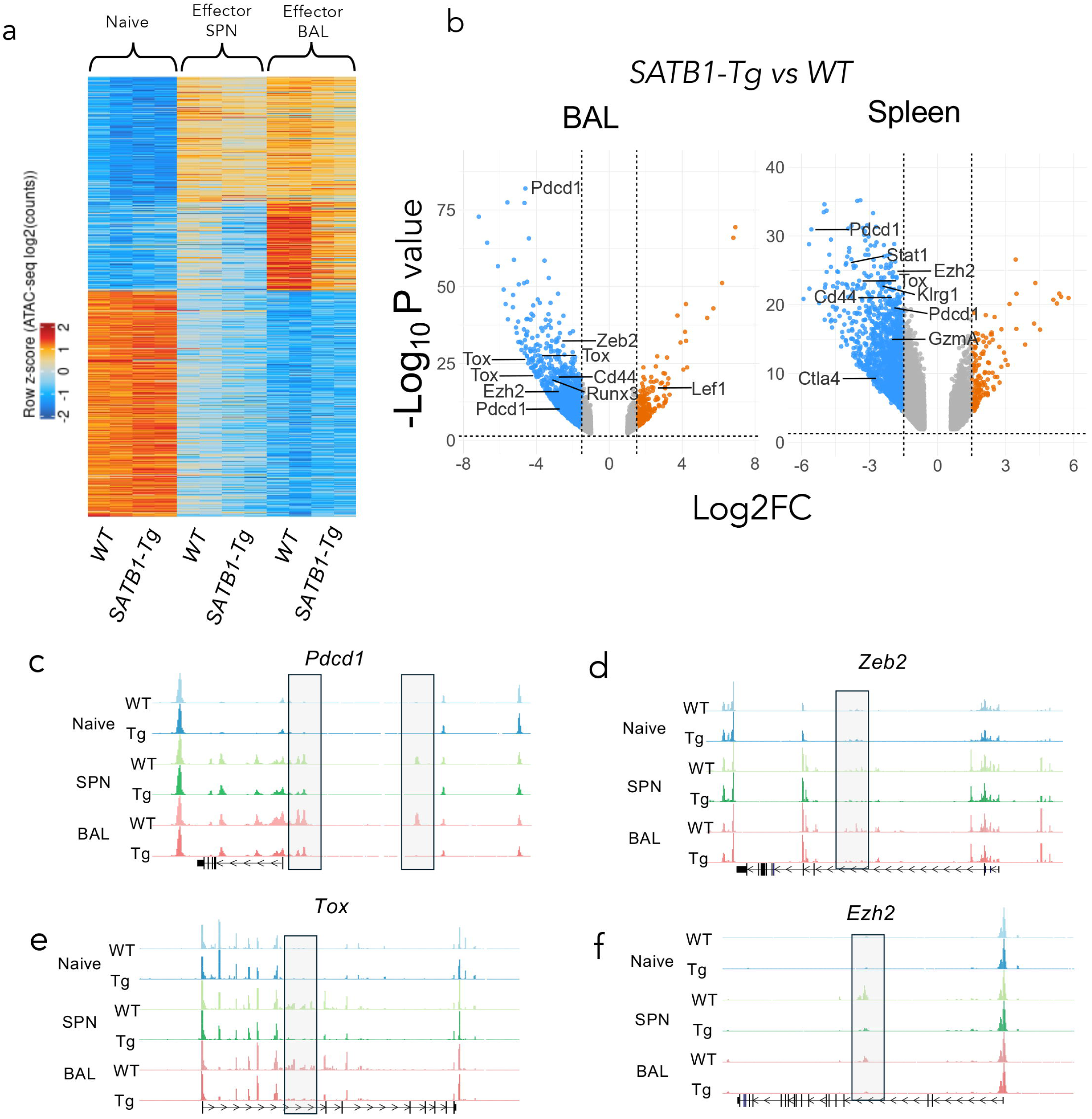
SATB1 overexpression limits chromatin accessibility at key CD8^+^ T cell effector gene loci. (**a**) ATAC-sequencing was carried out on sort purified naïve (CD44^lo^CD62L^hi^) or day 10 effector (BAL and Spleen) WT and SATB1-Tg OT-1 cells. ATAC-sequencing data was processed and *k*-means clustering carried out on differentially accessible regions (DARs) identified across the distinct CD8^+^ T cell subsets, selected with a log_10_ false discovery rate (FDR) of <0.01 and absolute log_2_ fold change (logFC) of >3). Shown is a heat map generated based on *z*-score. (**b**) Volcano plots of DARs linked to nearest neighbouring gene transcriptional start sites filtered based on a Log_10_FDR <0.1 and log_2_FC >1.5 across effector BAL and splenic effector OT-1 subsets comparing accessibility of SATB1-Tg relative to WT. (c-f) ATAC-sequencing tracks of naïve and effector (BAL and spleen) WT and SATB1-Tg OT-1 CD8+ T cells showing *Pdcd1* (**c**), *Zeb2* (**d**), *Tox* (**e**) and *Ezh2* (**f**) gene loci.

To further dissect how SATB1 binding can limit CD8^+^ T cell differentiation, we assessed enrichment of transcription factor binding sites (TFBS) at DARs identified between WT and SATB1-Tg effector CD8^+^ T cells (**Fig. 6**). We utilised the GIGGLE integration tool^49^ to assess the overlap of TF binding potential to DARs observed in SATB1-Tg vs WT effector CD8+ T cells from the BAL and spleen (**Fig. 6a**). Binding peaks for TFs such as RUNX3, members of the AP-1 complex (BATF, Fos, Jun) family, members of the STAT family (STAT4, STAT5, STAT6), IRF4, as well as the histone acetyltransferase EP300 were all found to overlap with DARs identified between SATB1-Tg and WT effector OT-I CD8^+^ T cells. Further, *de novo* motif enrichment analysis for DARs identified motifs that are targets for TFs such as Bcl11b, Runx3, BATF and SATB1 (**Fig. 6b**). Interesting, TF motifs for Elf1/3 and Znf348 (also known as ZBTB33) were also identified but have no known function in CD8+ T cell differentiation. Collectively, these data suggest that SATB1 regulates CD8^+^ T cell differentiation by limiting chromatin accessibility at transcriptional enhancers that are targets for TFs that would otherwise promote an effector transcriptional program.

**Figure 6.**
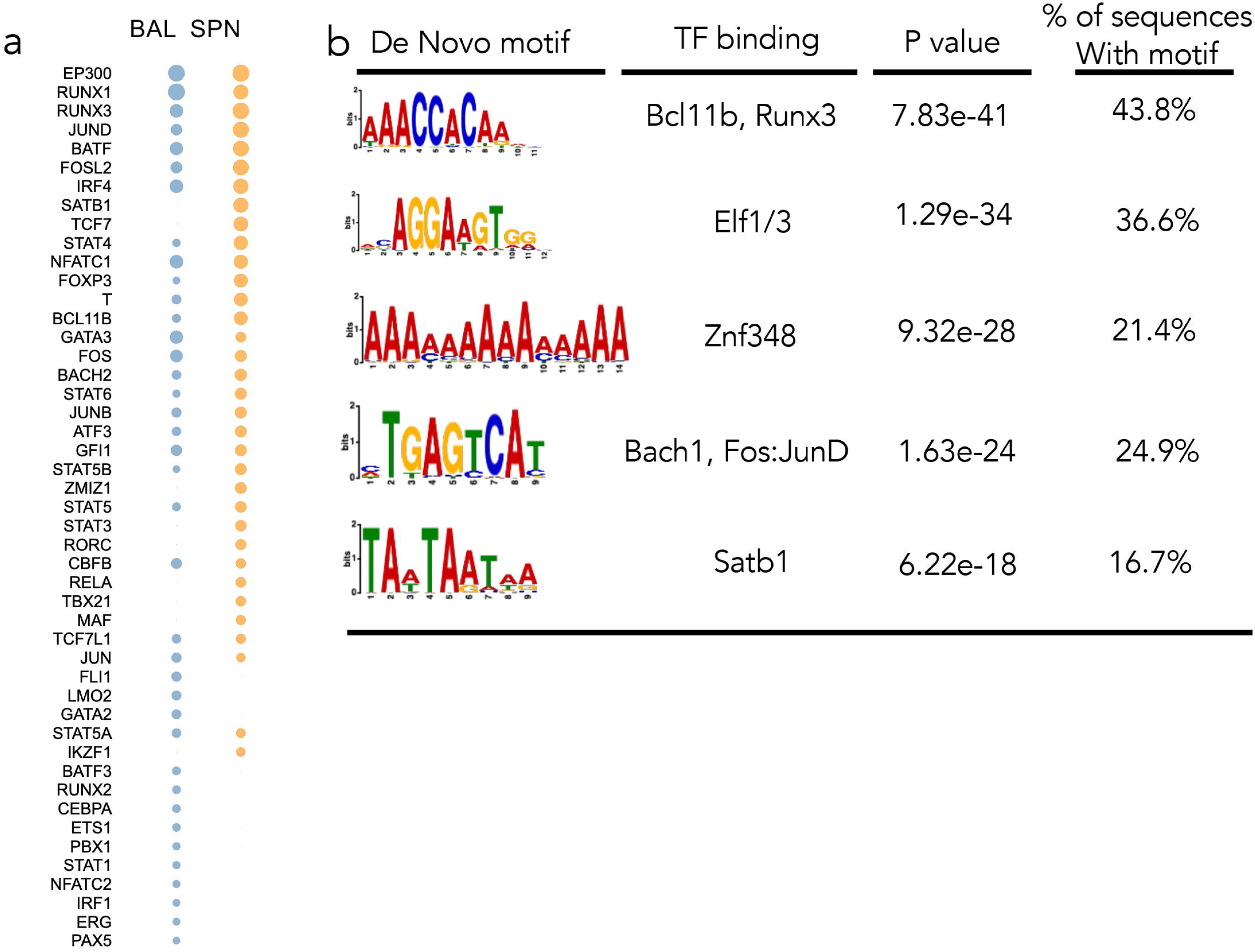
SATB1 overexpression limits accessibility of genomic regions targeted by effector associated transcription factors. (**a**) DARs found to be more accessible in either BAL or splenic effector WT vs SATB1-Tg OT-1 effector CD8^+^ T cells were identified and cross-referenced against publicly available ChIP-seq binding data using the Giggle tool^49^. TF binding sites enriched in DARs more accessible in BAL and spleen were listed according to significance. (**b**) *De novo* motif enrichment for specific TF binding sites in DARs was carried out using HOMER. Shown are those motifs that were most significantly enriched at DARs found between WT and SATB1-Tg effector CD8^+^ OT-I T cells.

## Discussion

The transition of CD8^+^ T cells from a naïve to effector state upon activation is normally associated with a link between extended proliferation and acquisition of signature phenotypic and functional markers. There is extensive literature identifying TFs that are induced upon CD8^+^ T cell activation, and how they then regulate the active process of effector differentiation^21, 22, 23, 24, 50, 51, 52^. Conversely, TFs highly expressed in the naïve state are often downregulated upon activation to enable full effector differentiation^28, 30^. Importantly, the suite of naïve TFs is still relatively limited in scope. Integrated analysis of single cell RNA and ATAC-seq data in naïve and effector CD8 T cells identified SATB1 as highly expressed in naive T cells with drastic down regulation upon effector differentiation. Functionally, OT-I CD8^+^ T cells engineered to sustain SATB1 expression after activation, demonstrated lower levels of effector differentiation, and a more stem-like memory phenotype. The inability of activated SATB1-Tg OT-I cells to express markers of effector differentiation corresponded to limited chromatin accessibility at non-coding regulatory elements linked to these gene loci. Interestingly, while SATB1 over expression limited acquisition of effector gene expression, it did not impact cellular proliferation, suggesting that these two aspects of CD8^+^ T cell differentiation can be uncoupled. Given our data also demonstrated this was linked to a more inaccessible chromatin structure, this study identifies SATB1 as a key transcriptional repressor that contributes to maintaining the naïve state by limiting acquisition of signature transcriptional programs. Collectively, our data position SATB1 as a central gatekeeper of effector T cell differentiation, in a manner that is independent of cellular division.

SATB1-overexpression resulted in decreased PD-1 expression in IAV-specific and OT-I CD8^+^ T cells across all timepoints assessed. Down-regulation of SATB1 correlates with increased and stable PD-1 expression in lymphocytic choriomeningitis virus (LCMV) mediated CD8^+^ T cell exhaustion^53^. Intriguingly, SATB1 over expression did not alter chromatin accessibility at the *Pdcd1* TSS, but did decreased accessibility at a key transcriptional enhancer found-23Kb upstream of the TSS, that is specifically regulated during CD8^+^ T cell exhaustion ^46^. We have previously demonstrated that SATB1 binds non-coding regulatory elements linked to effector gene loci, and that perturbation of this binding in naïve T cells results in a premature effector transcriptional signature^27, 43^. These data, along with similar findings at other gene loci, suggest that while SATB1 expression has a profound impact on effector CD8^+^ T cell phenotype and function, its effect is via highly targeted regulation of discrete non-coding regulatory regions in the genome. In addition, there was also reduced chromatin accessibility at key regulatory elements associated with the *Tox* locus, also shown to be a key driver of T cell exhaustion^47, 48^. Certainly, recent data from Kallies and colleagues demonstrated that SATB1 deficiency increases the generation of exhausted T cells after chronic LCMV infection, while CD8^+^ T cell specific SATB1 overexpression promotes the generation of TPEX during chronic LCMV infection (data not shown, personnel communication). Given TPEX are progenitor CD8^+^ T cells that respond to immune checkpoint blockade^54, 55^, targeting SATB1 to promote expression may therefore have therapeutic implications for treatment of chronic-related diseases and cancer.

A key observation was that while the acquisition of differentiation markers was impacted by SATB1 overexpression, the proliferative response of virus-specific CD8^+^ T cells was not overly impacted, particularly in response to secondary infection. It has long been suggested that the acquisition of effector function (differentiation) is intrinsically linked to proliferation^1, 3, 9, 17, 18, 19, 56^. However, our data suggest that proliferative capacity and effector CD8^+^ T cell differentiation can be decoupled, and that regulation of differentiation can occur directly, and not as a secondary consequence of controlling T cell activation or proliferation. The generation of effector CD8^+^ T cell responses often comes at the expense of generating optimal T cell memory ^1^. Given that SATB1 is in fact initially upregulated after T cell activation, this suggests that SATB1 may act as an early checkpoint that limits acquisition of effector phenotype and function early after activation. As such, SATB1 likely contributes to promoting the establishment of a pool of memory precursors early by limiting the transition to a mature effector state, while not impeding the early proliferative response.

We have previously shown that SATB1 binding is enriched at effector gene loci in naïve CD8^+^ T cells ^43^. The current study used integrated single-cell genomic analyses to identify SATB1 as a candidate transcriptional repressor; a notion that was verified by SATB1 overexpression reducing the chromatin accessibility within effector CD8^+^ T cells, thus providing mechanistic insight into SATB1 function. Of particular interest was our observation that genome regions that remained more inaccessible due to SATB1 overexpression were enriched for binding sites for TFs associated with effector T cell differentiation. This included TFBSs for Activator Protein-1 (AP-1) TF family members JUN (JUNB, JUND), FOS, NFATc1, BATF, IRF4 and a variety of STAT family members (STAT4, STAT5A, STAT5B). The role of SATB1 in establishing a transcriptionally repressive chromatin landscape may be in part due to recruitment of HDACs to regulatory elements, as described for the *Pdcd1* TSS in tumour-specific CD8^+^ T cells ^57^. In this way, SATB1 limits binding of effector TFs by limiting the necessary chromatin remodelling to expose TFBS at regulatory elements. Interestingly, BACH2 TFBSs were also enriched in WT effector CD8^+^ T cells compared to SATB1-Tg CD8+ T cells. BACH2 limits effector differentiation by occupying AP1 binding sites in naïve CD8^+^ T cells and needs to be downregulated upon activation, thus exposing those regions for AP-1 targeting ^33, 34^. We have demonstrated that BACH2 and SATB1 act as regulators of higher order chromatin structures in naïve CD8^+^ T cells, and that loss of SATB1 and BACH2 binding in naïve CD8^+^ T cells results in chromatin remodelling that mirrors that found in effector CD8^+^ T cells ^27^. Hence, it is also likely that SATB1 collaborates with BACH2 to occupy key regulatory sites that would otherwise be targeted by effector TFs. Together, these data highlight how SATB1 serves to maintain T cell naïvety and limit effector differentiation early after T cell activation to promote memory formation.

## Acknowledgements

We thank members of the Turner and La Gruta labs for helpful discussions and critical reading of the manuscript. We thank the Monash Flowcore facility (Monash University, Clayton) for helpful advice and technical assistance with flow cytometry and cell sorting experiments, and the Monash Micromon Genomics (Monash University, Clayton) and Hudson Genomics (Hudson Institute, Monash University Health and Medical Precinct, Clayton) facilities for advice relating RNA-Seq. We thank the LIMES-GRC for the generation of SATB-Tg mice.

Mice were bred and housed at the Monash Animal Research Platform, at Monash University, Clayton. This work was supported by grants from the National Health and Medical Research Council of Australia, APP1003131 (S.J.T); an Australian Research Council Discovery Grant, DP170102020 (S.J.T); Australian Postgraduate Awards (J.K.C.L; T.J.B.); MDB is supported by the Helmholtz Association and the German Research Foundation (DFG) (SFB1454 project number 432325352, IGK2168/2 project number 272482170). MDB and JLS are members of the excellence cluster ImmunoSensation2 (EXC2151 project number 390873048) and the EU Horizon 2020 project SYSCID under grant agreement no. 733100. JLS received funds from the DFG (IGK2168/2 project number 272482170) and the BMBF (DietBB - 01EA1809A, iTREAT - 01ZX1902A).

## Materials and methods

### Mice and co-adoptive transfer

WT, WT OT-I, *SATB1*-*GFP* x *Lck^CRE^* (SATB1-Tg) and *SATB1*-*GFP* x *Lck^CRE^* x OT-I (SATB1-Tg OT-I) with congenic markers (CD45.1^+^, CD45.1/45.2^+^ or CD45.2^+^) are C57BL/6 (B6, H-2D^b^/K^b^) mice that were bred by the Monash Animal Research Platform, Clayton. SATB1-Tg and SATB1-Tg OT-1 mice expressing GFP^+^ CD8^+^ T cells were identified by flow cytometry and genotyping. For co-adoptive transfers, naïve (CD8^+^ CD44^low^) WT OT-I cells (CD45.1^+^) and SATB1-Tg OT-I cells (CD45.1/45.2^+^) were FACS purified and pooled, and 1 × 10^4^ of each were adoptively transferred by retro-orbital injection into naïve B6 (CD45.2^+^) recipient mice, 24-hours prior to IAV infection. All animal experiments were approved by the Monash Animal Ethics Committee.

### Influenza A Virus (IAV)infection

For primary IAV infections, A/HKx31 (H3N2) was used to infect WT B6 and SATB1-Tg B6 mice, and A/HKx31-OVA was used to initiate WT and SATB1-Tg OT-I T cell responses. In each case, 1 x 10^4^ pfu was delivered intranasally. For secondary infections, mice primed as above were rechallenged with 1 x 10^3^ pfu of A/PR8 (H1N1) or A/PR8-OVA at 60 days post-primary infection.

### Flow cytometry

Prior to surface staining, lymphocyte preparations were treated with 2.4G2 Fc receptor block. For intracellular cytokine staining, cells from WT/SATB1-Tg were restimulated with 1μM NP_366-374_/PA_224-233_, or N4 (SIINFEKL, OVA_257-264_), in the presence of 10U/mL rhIL-2 and GolgiPlug diluted in complete RPMI, for 5 hours. Cells were then fixed and permeabilized with Cytofix/Cytoperm (BD Bioscience) kit according to manufacturer’s instructions. For intranuclear staining of TFs, cells were fixed and permeabilized with FoxP3/Transcription Factor Staining Buffer set (eBioscience). Fixable Live/Dead viability dye was used to determine live populations. Cell populations were characterised using BD Fortessa or BD FACSymphony A3 (Monash FlowCore Facility) and analysed with FlowJo version 10 software.

### RNA-sequencing

Sort-purified naïve and effector OT-I cells were lysed with TRIzol followed by RNA extraction using Direct-zol^TM^ RNA Miniprep Plus kit (Zymo Research). Preparation of samples for RNA-sequencing was performed according to Russ et al. (2014)^58^, and sequenced with an Illumina NextSeq 2000 by the Monash Health Translational Precinct (MHTP) Medical Genomics Facility at the Hudson Institute of Medical Research. Data analysis was performed by the Monash Bioinformatics Platform.

### Assay for transposase-accessible chromatin with sequencing (ATAC-seq)

The ATAC-seq protocol performed here was adapted from Buenrostro et al. (2015)^59^. Nuclei from 5 x 10^4^ OT-I cells were isolated using cold lysis buffer. Nuclei were resuspended in Tn5 reaction mix and incubated at 37°C for 30 minutes for transposition before tagmented DNA was purified using the MinElute PCR Purification kit (Qiagen). Libraries were apmplified with 5 PCR cycles using i5 index PCR primers described in Buenrostro et al. (2013)^60^. Partially amplified libraries were then subjected to 20 cycles in a real-time quantitative PCR to determine the optimal number of PCR cycles needed for library amplification. Amplified DNA was purified using the MinElute PCR Purification kit (Qiagen). Libraries were subjected to paired –end seuqencing on an Illumina NextSeq 2000 as above.

### Single cell data analysis

The single cell RNAseq data was processed using the cellranger pipeline (cellranger version 3.0.2, cellranger mm10 reference: refdata-cellranger-mm10-3.0.0), converting the raw fastq files to count matrices for further analysis. The individual samples were then processed using Seurat (v4.0.1) and scater (v1.18.6). Quality control was performed to filter out cells with low number of genes detected as detected by the isOutlier function from scater, batched by sample id and log transformed. Additionally, outlier cells for both high and low library size were filtered out as well as cells with high percentage of mitochondrial reads. After cell filtering, Seurat was used for log transformation and normalisation. Dimensionality reduction was performed with standard Seurat parameters to generate an UMAP.

The single cell ATACseq data was processed using the cellranger atac pipeline (cellranger version 2.0.0, cellranger mm10 reference: refdata-cellranger-arc-mm10-2020-A-2.0.0). The individual samples were read into Signac (v1.5.0) for analysis. Outlier cells were identified using the isOutlier function using scuttle (v1.0.4) to detect cells with low and high numbers of peak counts, low numbers of detected peaks. These cells were removed along with cells that failed to have at least 20 percent of reads in called peaks, a nucleosome signal > 4 and a TSS score < 2. Once the low quality cells had been filtered out, a consensus peak set was called again with MACS2 on a sample basis with the ‘reduce’ stragy via Signac’s CallPeaks function. A new peak count matrix was generated and this peak set was used for all downstream analysis with Seurat. Harmony (v1.0) was used to batch correct samples, as the naive, effector_1 and memory samples were generated in a separate batch to effector_2, bal_1 and bal_2 samples. Seurat’s FindTransferAnchors function was used to transfer cell type labels from the scRNA dataset to the scATAC dataset.

For SCENIC+ analysis (v0.1.dev455), the scATAC dataset was re-analysed using pycisTopic and pycistarget for quality control, topic modelling, calculating differentially accessible regions and genes and motif enrichment. A custom motif database was generated using the create cisTarget databases tool. As input, it was provided the consensus peaks generated by pycisTopic and the custom motif database was used as input for the motif enrichment testing - using a custom motif database enabled the calculation of two Satb1 eRegulons instead of one when the prebuilt mm10 motif database was used. The RNA samples were converted from Seurat to Scanpy to avoid re-processing the data. The SCENIC workflow was followed to calculate eRegulons from the differentially expressed genes, differentially accessible regions/genes and the differentially enriched motifs. The eRegulons were not filtered after calculation in order to examine the SATB1 eRegulons generated by the dataset. Additionally, perturbation simulations were performed to simulate how knockdown and overexpression of Satb1 would affect cell type differentiation.

## References

1. Kaech, S.M. & Ahmed, R. Memory CD8+ T cell differentiation: initial antigen encounter triggers a developmental program in naive cells. Nat Immunol 2, 415–422 (2001).

2. van Stipdonk, M.J. et al. Dynamic programming of CD8+ T lymphocyte responses. Nat Immunol 4, 361–365 (2003).

3. Kaech, S.M., Hemby, S., Kersh, E. & Ahmed, R. Molecular and functional profiling of memory CD8 T cell differentiation. Cell 111, 837–851 (2002).

4. Lalvani, A. et al. Rapid effector function in CD8+ memory T cells. J Exp Med 186, 859–865 (1997).

5. Oehen, S. & Brduscha-Riem, K. Differentiation of naive CTL to effector and memory CTL: correlation of effector function with phenotype and cell division. J Immunol 161, 5338–5346 (1998).

6. Jenkins, M.R., Kedzierska, K., Doherty, P.C. & Turner, S.J. Heterogeneity of effector phenotype for acute phase and memory influenza A virus-specific CTL. J Immunol 179, 64–70 (2007).

7. Nguyen, M.L. et al. Dynamic regulation of permissive histone modifications and GATA3 binding underpin acquisition of granzyme A expression by virus-specific CD8(+) T cells. Eur J Immunol 46, 307–318 (2016).

8. Peters, P.J. et al. Cytotoxic T lymphocyte granules are secretory lysosomes, containing both perforin and granzymes. J Exp Med 173, 1099–1109 (1991).

9. Denton, A.E., Russ, B.E., Doherty, P.C., Rao, S. & Turner, S.J. Differentiation-dependent functional and epigenetic landscapes for cytokine genes in virus-specific CD8+ T cells. Proceedings of the National Academy of Sciences of the United States of America 108, 15306–15311 (2011).

10. La Gruta, N.L., Turner, S.J. & Doherty, P.C. Hierarchies in cytokine expression profiles for acute and resolving influenza virus-specific CD8+ T cell responses: correlation of cytokine profile and TCR avidity. J Immunol 172, 5553–5560 (2004).

11. Slifka, M.K., Rodriguez, F. & Whitton, J.L. Rapid on/off cycling of cytokine production by virus-specific CD8+ T cells. Nature 401, 76–79 (1999).

12. Russ, B.E. et al. Regulation of H3K4me3 at Transcriptional Enhancers Characterizes Acquisition of Virus-Specific CD8(+) T Cell-Lineage-Specific Function. Cell Rep 21, 3624–3636 (2017).

13. Schall, T.J. et al. A human T cell-specific molecule is a member of a new gene family. J Immunol 141, 1018–1025 (1988).

14. Barber, D.L. et al. Restoring function in exhausted CD8 T cells during chronic viral infection. Nature 439, 682–687 (2006).

15. Ishida, Y., Agata, Y., Shibahara, K. & Honjo, T. Induced expression of PD-1, a novel member of the immunoglobulin gene superfamily, upon programmed cell death. EMBO J 11, 3887–3895 (1992).

16. Kohlmeier, J.E. et al. Inflammatory chemokine receptors regulate CD8(+) T cell contraction and memory generation following infection. J Exp Med 208, 1621–1634 (2011).

17. Badovinac, V.P., Haring, J.S. & Harty, J.T. Initial T cell receptor transgenic cell precursor frequency dictates critical aspects of the CD8(+) T cell response to infection. Immunity 26, 827–841 (2007).

18. Lawrence, C.W. & Braciale, T.J. Activation, differentiation, and migration of naive virus-specific CD8+ T cells during pulmonary influenza virus infection. J Immunol 173, 1209–1218 (2004).

19. Moffat, J.M., Gebhardt, T., Doherty, P.C., Turner, S.J. & Mintern, J.D. Granzyme A expression reveals distinct cytolytic CTL subsets following influenza A virus infection. Eur J Immunol 39, 1203–1210 (2009).

20. Jenkins, M.R. et al. Cell cycle-related acquisition of cytotoxic mediators defines the progressive differentiation to effector status for virus-specific CD8+ T cells. J Immunol 181, 3818–3822 (2008).

21. Cruz-Guilloty, F. et al. Runx3 and T-box proteins cooperate to establish the transcriptional program of effector CTLs. J Exp Med 206, 51–59 (2009).

22. Intlekofer, A.M. et al. Effector and memory CD8+ T cell fate coupled by T-bet and eomesodermin. Nat Immunol 6, 1236–1244 (2005).

23. Kallies, A., Xin, A., Belz, G.T. & Nutt, S.L. Blimp-1 transcription factor is required for the differentiation of effector CD8(+) T cells and memory responses. Immunity 31, 283–295 (2009).

24. Man, K. et al. The transcription factor IRF4 is essential for TCR affinity-mediated metabolic programming and clonal expansion of T cells. Nat Immunol 14, 1155–1165 (2013).

25. Yao, S. et al. Interferon regulatory factor 4 sustains CD8(+) T cell expansion and effector differentiation. Immunity 39, 833–845 (2013).

26. Bennett, T.J., Udupa, V.A.V. & Turner, S.J. Running to Stand Still: Naive CD8(+) T Cells Actively Maintain a Program of Quiescence. Int J Mol Sci 21 (2020).

27. Russ, B.E., et al. Active maintenance of CD8(+) T cell naivety through regulation of global genome architecture. bioRxiv (2023).

28. Delpoux, A. et al. FOXO1 constrains activation and regulates senescence in CD8 T cells. Cell Rep 34, 108674 (2021).

29. Kerdiles, Y.M. et al. Foxo1 links homing and survival of naive T cells by regulating L-selectin, CCR7 and interleukin 7 receptor. Nat Immunol 10, 176–184 (2009).

30. Shan, Q. et al. Tcf1 and Lef1 provide constant supervision to mature CD8(+) T cell identity and function by organizing genomic architecture. Nat Commun 12, 5863 (2021).

31. Gray, S.M., Amezquita, R.A., Guan, T., Kleinstein, S.H. & Kaech, S.M. Polycomb Repressive Complex 2-Mediated Chromatin Repression Guides Effector CD8(+) T Cell Terminal Differentiation and Loss of Multipotency. Immunity 46, 596–608 (2017).

32. Kakaradov, B. et al. Early transcriptional and epigenetic regulation of CD8(+) T cell differentiation revealed by single-cell RNA sequencing. Nat Immunol 18, 422–432 (2017).

33. Roychoudhuri, R. et al. BACH2 regulates CD8(+) T cell differentiation by controlling access of AP-1 factors to enhancers. Nat Immunol 17, 851–860 (2016).

34. Yao, C. et al. BACH2 enforces the transcriptional and epigenetic programs of stem-like CD8(+) T cells. Nat Immunol 22, 370–380 (2021).

35. Jenkins, M.R., Webby, R., Doherty, P.C. & Turner, S.J. Addition of a prominent epitope affects influenza A virus-specific CD8+ T cell immunodominance hierarchies when antigen is limiting. J Immunol 177, 2917–2925 (2006).

36. Russ, B.E. et al. Distinct Epigenetic Signatures Delineate Transcriptional Programs during Virus-Specific CD8(+) T Cell Differentiation. Immunity 41, 853–865 (2014).

37. Scott-Browne, J.P. et al. Dynamic Changes in Chromatin Accessibility Occur in CD8+ T Cells Responding to Viral Infection. Immunity 45, 1327–1340 (2016).

38. Yu, B. et al. Epigenetic landscapes reveal transcription factors that regulate CD8(+) T cell differentiation. Nat Immunol 18, 573–582 (2017).

39. Bravo Gonzalez-Blas, C., et al. SCENIC+: single-cell multiomic inference of enhancers and gene regulatory networks. Nat Methods 20, 1355–1367 (2023).

40. Wang, D. et al. The Transcription Factor Runx3 Establishes Chromatin Accessibility of cis-Regulatory Landscapes that Drive Memory Cytotoxic T Lymphocyte Formation. Immunity 48, 659–674 e656 (2018).

41. Olesin, E., Nayar, R., Saikumar-Lakshmi, P. & Berg, L.J. The Transcription Factor Runx2 Is Required for Long-Term Persistence of Antiviral CD8(+) Memory T Cells. Immunohorizons 2, 251–261 (2018).

42. Guan, T. et al. ZEB1, ZEB2, and the miR-200 family form a counterregulatory network to regulate CD8(+) T cell fates. J Exp Med 215, 1153–1168 (2018).

43. Nussing, S. et al. SATB1 ensures appropriate transcriptional programs within naive CD8(+) T cells. Immunol Cell Biol 100, 636–652 (2022).

44. Nussing, S. et al. Divergent SATB1 expression across human life span and tissue compartments. Immunol Cell Biol 97, 498–511 (2019).

45. Kohne, M. et al. Satb1 directs the differentiation of T(H)17 cells through suppression of IL-2 expression. Cell Rep 44, 115866 (2025).

46. Sen, D.R. et al. The epigenetic landscape of T cell exhaustion. Science 354, 1165–1169 (2016).

47. Khan, O. et al. TOX transcriptionally and epigenetically programs CD8(+) T cell exhaustion. Nature 571, 211–218 (2019).

48. Yao, C. et al. Single-cell RNA-seq reveals TOX as a key regulator of CD8(+) T cell persistence in chronic infection. Nat Immunol 20, 890–901 (2019).

49. Layer, R.M. et al. GIGGLE: a search engine for large-scale integrated genome analysis. Nat Methods 15, 123–126 (2018).

50. Kurachi, M. et al. The transcription factor BATF operates as an essential differentiation checkpoint in early effector CD8+ T cells. Nat Immunol 15, 373–383 (2014).

51. Prier, J.E. et al. Early T-BET Expression Ensures an Appropriate CD8(+) Lineage-Specific Transcriptional Landscape after Influenza A Virus Infection. J Immunol 203, 1044–1054 (2019).

52. Tsao, H.W., et al. Batf-mediated epigenetic control of effector CD8(+) T cell differentiation. Sci Immunol 7, eabi4919 (2022).

53. Wherry, E.J. et al. Molecular signature of CD8+ T cell exhaustion during chronic viral infection. Immunity 27, 670–684 (2007).

54. Utzschneider, D.T. et al. T Cell Factor 1-Expressing Memory-like CD8(+) T Cells Sustain the Immune Response to Chronic Viral Infections. Immunity 45, 415–427 (2016).

55. Utzschneider, D.T. et al. Early precursor T cells establish and propagate T cell exhaustion in chronic infection. Nat Immunol 21, 1256–1266 (2020).

56. Mercado, R. et al. Early programming of T cell populations responding to bacterial infection. J Immunol 165, 6833–6839 (2000).

57. Stephen, T.L. et al. SATB1 Expression Governs Epigenetic Repression of PD-1 in Tumor-Reactive T Cells. Immunity 46, 51–64 (2017).

58. Russ, Brendan E. et al. Distinct Epigenetic Signatures Delineate Transcriptional Programs during Virus-Specific CD8+ T Cell Differentiation. Immunity 41, 853–865 (2014).

59. Buenrostro, J.D., Wu, B., Chang, H.Y. & Greenleaf, W.J. ATAC-seq: A Method for Assaying Chromatin Accessibility Genome-Wide. Curr Protoc Mol Biol 109, 21.29.21–21.29.29 (2015).

60. Buenrostro, J.D., Giresi, P.G., Zaba, L.C., Chang, H.Y. & Greenleaf, W.J. Transposition of native chromatin for fast and sensitive epigenomic profiling of open chromatin, DNA-binding proteins and nucleosome position. Nature Methods 10, 1213–1218 (2013).

