## Supplemental Figures 1-6 for "SATB1 maintains naive-like identity in antiviral CD8□ T cells by limiting chromatin remodelling at effector gene loci"

a

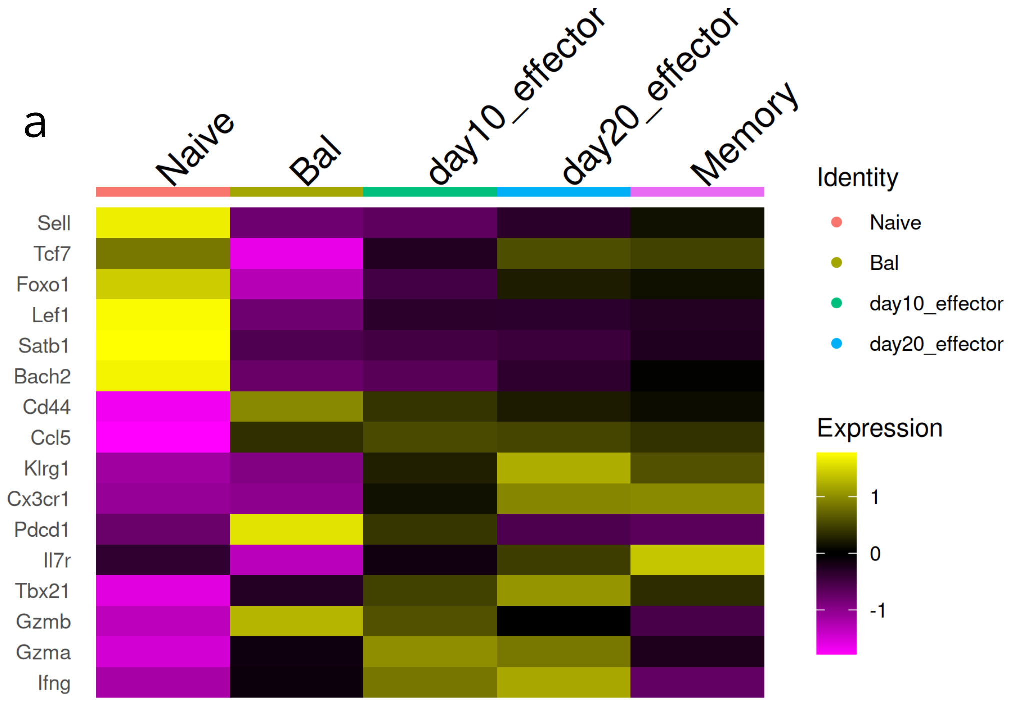
Naive
Bal
day10_effector
day20_effector
Memory
Sell
Tcf7
Foxo1
Lef1
Satb1
Bach2
Cd44
Ccl5
Klrg1
Cx3cr1
Pdcd1
Il7r
Tbx21
Gzmb
Gzma
Ifng
Identity
Naive
Bal
day10_effector
day20_effector
Expression
1
0
-1

b

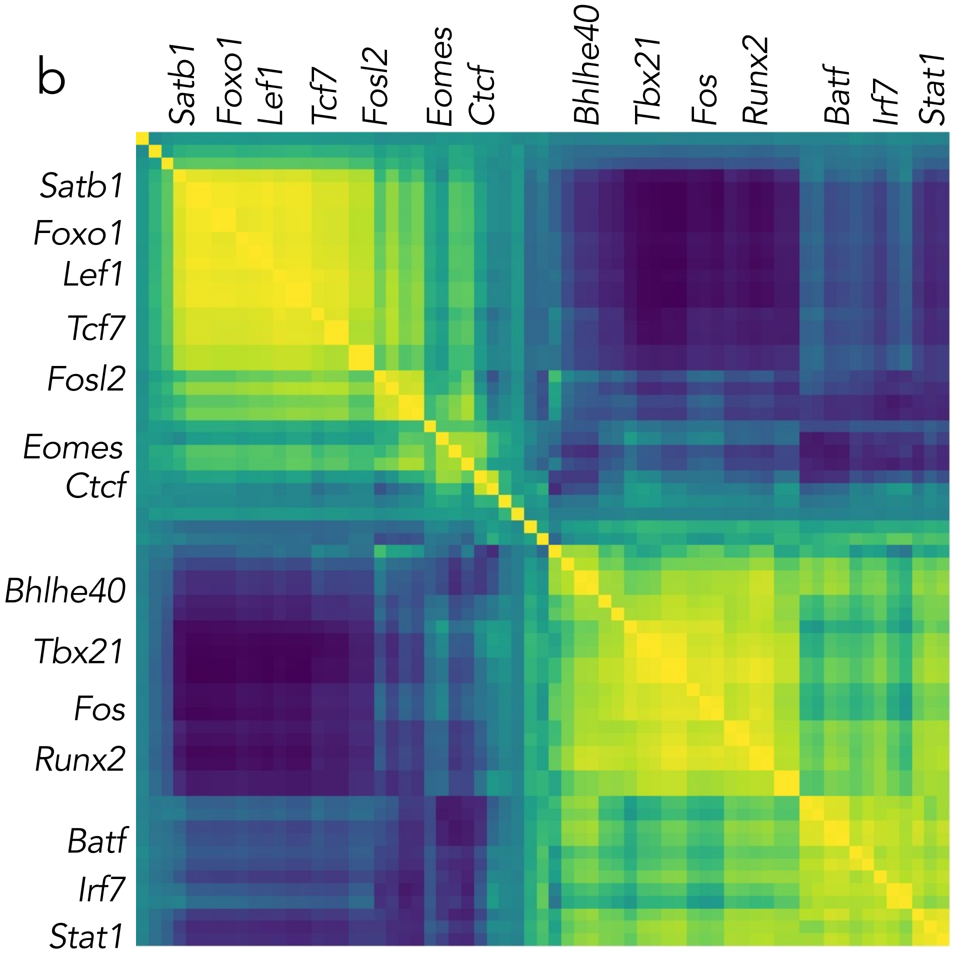
Satb1
Foxo1
Lef1
Tcf7
Fosl2
Eomes
Ctcf
Bhlhe40
Tbx21
Fos
Runx2
Batf
Irf7
Stat1

**Supplementary Figure 1. (a) Single cell RNA-seq mapping of key CD8$^+$ T cell subset signatures.** The transcripts for genes characteristic of naïve, effector and memory CD8$^+$ T differentiation states were aggregated from single RNA-seq data isolated from adoptively transferred OT-Is isolated on Day 0 (naïve), day 10 (Effector, BAL and Spleen), day 20 (later effector/early memory) and day 90 (memory) after A/HKx31-OVA infection. Aggregated data was clustered based on enrichment for the genes listed and represented as a heatmap. (**b**) Correlation of eRegulons within CD8+ T cell subsets. SCENIC+, was used to identify enriched GRNs within cells and pairwise Pearson correlations were calculated between eRegulons to assess co-regulatory relationships. The resulting matrix highlights clusters of eRegulons with coordinated activity.

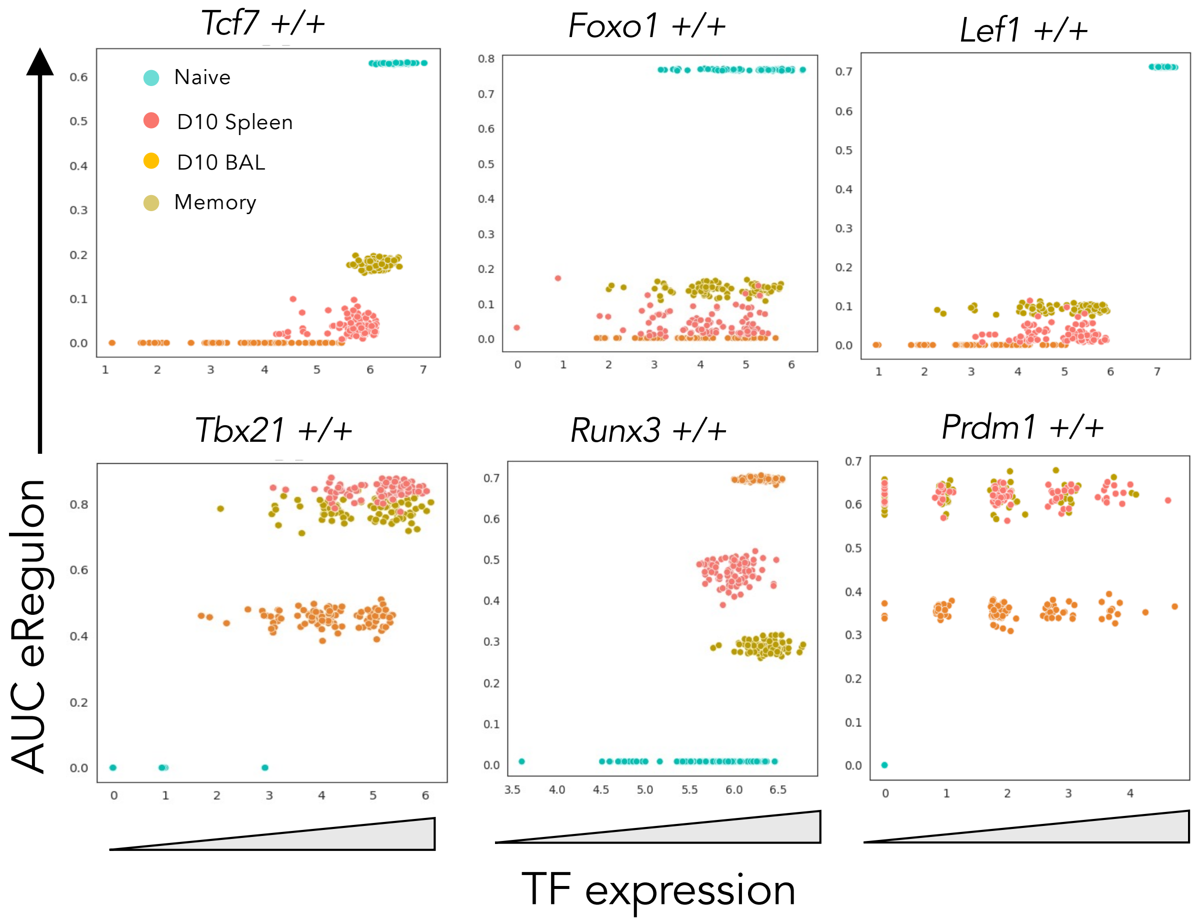


**Supplementary Figure 2. Assessment of SATB1 expression and the correlation with GRNs that exhibit accessible chromatin enriched with specific TF binding sites.** The plots show the level of expression of TFs known to be associated with the naïve state (TCF1, encoded by Tcf7; Foxo1; Lef1) or the effector state (Tbx21, Runx3 and Prdm1 encoding for BLIMP1) plotted against GRNs with open chromatin targeted by each TF and that exhibit positive gene regulation (+/+).

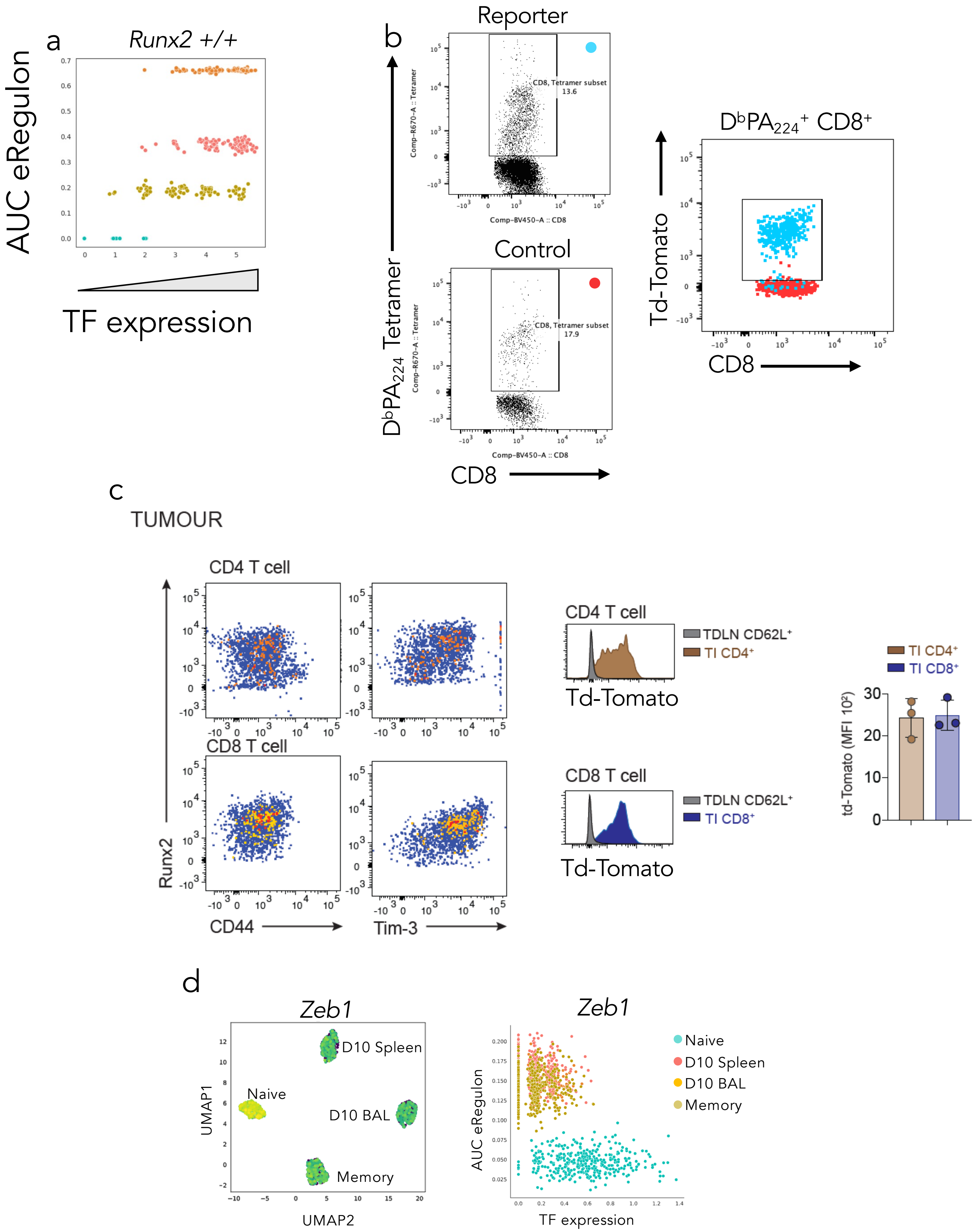
a
Runx2 +/+
AUC eRegulon
TF expression
b
Reporter
CD8, Tetramer subset 13.6
Control
CD8, Tetramer subset 17.9
Comp-R670-A :: Tetramer
Comp-BV450-A :: CD8
DbPA224 Tetramer
CD8
DbPA224+ CD8+
Td-Tomato
CD8
c
TUMOUR
CD4 T cell
CD8 T cell
Runx2
CD44
Tim-3
CD4 T cell
TDLN CD62L+
TI CD4+
Td-Tomato
CD8 T cell
TDLN CD62L+
TI CD8+
Td-Tomato
TI CD4+
TI CD8+
td-Tomato (MFI 10²)
d
Zeb1
D10 Spleen
Naive
D10 BAL
Memory
UMAP1
UMAP2
Zeb1
AUC eRegulon
TF expression
Naive
D10 Spleen
D10 BAL
Memory

**Supplementary Figure 3.** (**a**) Assessment of RUNX2 expression and correlation with GRNs identified in single cells from naïve, effector and memory OT-I T cell subsets. (**b, c**) Assessment of RUNX2 expression using RUNX2 reporter mice that have TdTomato knocked into *Runx2* locus. (**b**) WT or RUNX2 reporter mice were infected with A/HKx31 and effector $D^bPA_{224}$-specific $CD8^+$ T cells assessed for TdTomato expression by flow cytometry. (**c**) WT or RUNX2 reporter mice were challenged intradermally with MC38 tumours. Tumours were isolated 2 weeks after challenge and intratumoral CD8 and CD4 lymphocytes isolated and assessed for tdTomato expression by flow cytometry. (**d**) PCA based on GRN enrichment within naïve, effector and memory OT-I cells showing ZEB1 expression within each GRN PCA cluster. Assessment of ZEB1 expression and correlation with GRNs identified in single cells from naïve, effector and memory OT-I T cell subsets.

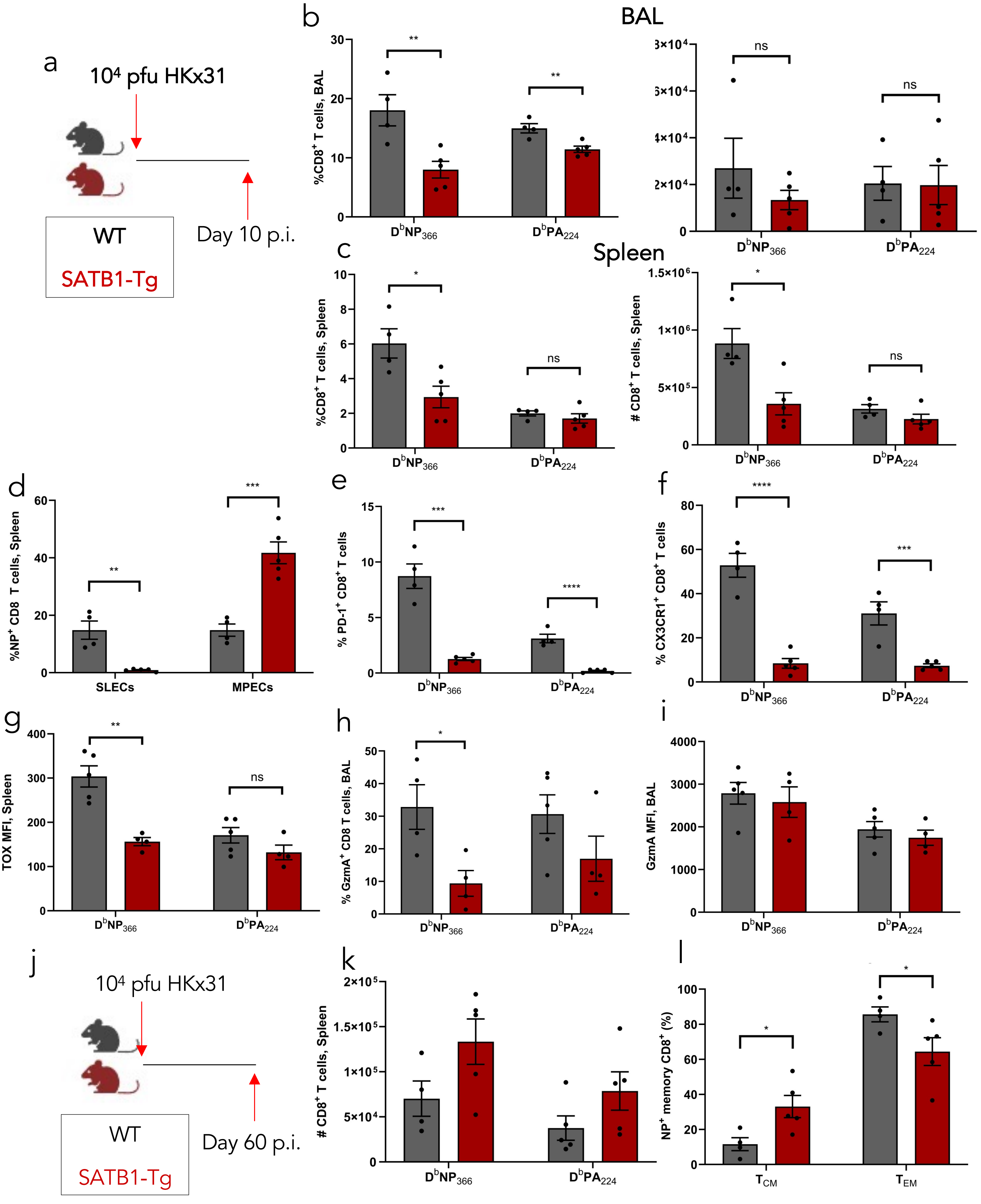
a
10⁴ pfu HKx31
WT
SATB1-Tg
Day 10 p.i.
b
BAL
%CD8⁺ T cells, BAL
**
**
ns
ns
DbNP366
DbPA224
c
Spleen
%CD8⁺ T cells, Spleen
### CD8⁺ T cells, Spleen
*
*
ns
ns
d
%NP⁺ CD8 T cells, Spleen
**
***
SLECs
MPECs
e
% PD-1⁺ CD8⁺ T cells
***
****
f
% CX3CR1⁺ CD8⁺ T cells
****
***
g
TOX MFI, Spleen
**
ns
h
% GzmA⁺ CD8 T cells, BAL
*
i
GzmA MFI, BAL
j
10⁴ pfu HKx31
WT
SATB1-Tg
Day 60 p.i.
k
### CD8⁺ T cells, Spleen
l
NP⁺ memory CD8⁺ (%)
*
*
TCM
TEM

**Supplementary Figure 4. SATB1 overexpression limits IAV-specific CD8+ T cell effector differentiation.** (**a**) Timeline for A/HKx31 infection of WT and SATB1-Tg mice. (**b**) On day 10 after primary A/HKx31 infection, the proportion and absolute numbers of $D^bNP_{366-374}$- and $D^bPA_{224-233}$-specific $CD8^+$ T cells were assessed from the BAL (**b**) and spleen (**c**). (**d-h**) Mean proportions of splenic $D^bNP_{366}$- and $D^bPA_{224}$-specific CD8 T cells that were defined as (**d**) SLECs ($KLRG1^{hi}CD127^{lo}$) or MPECs ($KLRG1^{lo}CD127^{hi}$) in WT and SATB1-Tg mice; (**e**) the proportion of $D^bNP_{366}$-specific $CD8^+$ T cells expressing PD-1; (**f**) CX3CR1; (**g**) the MFI of TOX and the proportion (**h**) and MFI (**i**) of Granzyme A (GzmA). (**j**) Timeline for A/HKx31 infection of WT and SATB1-Tg mice to look at memory $CD8^+$ T cells. (**k**) The number of $D^bNP_{366}$- and $D^bPA_{224}$-specific memory CD8+ T cells isolated at day 90 after primary A/HKx31 infection. (**i**) Mean proportions of splenic $D^bNP_{366}$-specific CD8 T cells defined as central memory ($T_{CM}$, $CD44^{hi}CD62L^{hi}$) or effector memory ($T_{EM}$, $CD44^{hi}CD62L^{hi}$) in WT and SATB1-Tg mice. Data is representative of three independent experimental repeats, N=4 female C57BL/6 mice and N=5 female *SATB1-GFP Lck*$^{Cre}$ (SATB1-Tg) mice. Data represented as mean ± SEM. * indicates $p$-value≤0.05, ** $p$≤0.01, *** $p$≤0.001 and *** $p$≤0.0001, unpaired student's $t$-test.

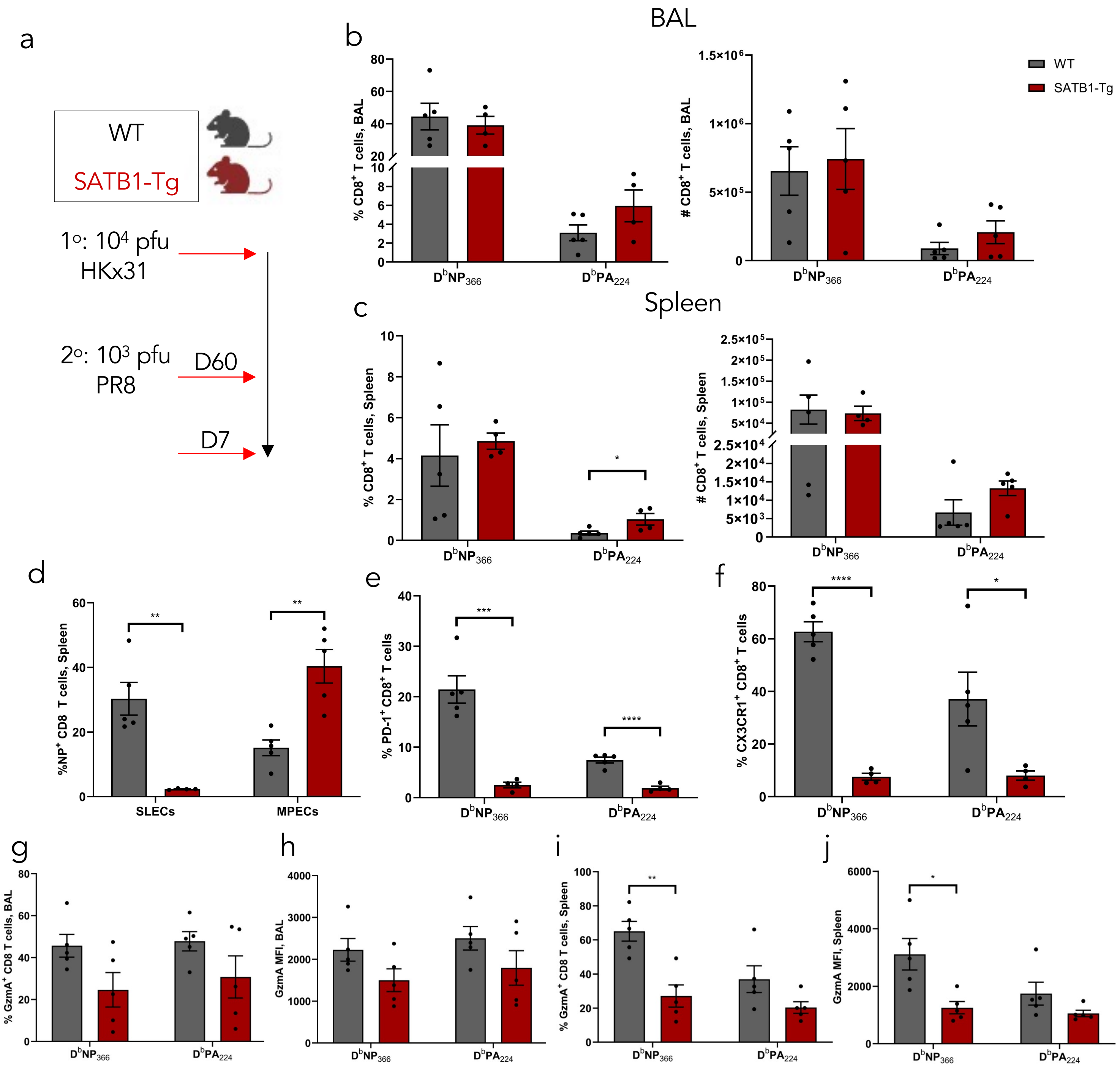
BAL
Spleen
a
WT
SATB1-Tg
1°: 10⁴ pfu
HKx31
2°: 10³ pfu
PR8
D60
D7
b
% CD8⁺ T cells, BAL
### CD8⁺ T cells, BAL
WT
SATB1-Tg
D^bNP366
D^bPA224
c
% CD8⁺ T cells, Spleen
### CD8⁺ T cells, Spleen
d
%NP⁺ CD8 T cells, Spleen
SLECs
MPECs
e
% PD-1⁺ CD8⁺ T cells
f
% CX3CR1⁺ CD8⁺ T cells
g
% GzmA⁺ CD8 T cells, BAL
h
GzmA MFI, BAL
i
% GzmA⁺ CD8 T cells, Spleen
j
GzmA MFI, Spleen

**Supplementary Figure 5.** (**a**) Timeline for A/PR8 (H1N1) infection of A/HKx31 (H3N2) primed WT and SATB1-Tg mice for analysis of secondary responses. (**b, c**) On day 7 after secondary A/PR8 infection of A/HKx31 primed mice, the proportion and absolute numbers of $D^bNP_{366\text{-}374}$- and $D^bPA_{224\text{-}233}$-specific $CD8^+$ T cells were assessed from the BAL (**b**) and spleen (**c**) of WT and SATB1-Tg mice. (**d**) Mean proportions of splenic $D^bNP_{366}$-specific CD8 T cells defined as (**d**) SLECs ($KLRG1^{hi}CD127^{lo}$) or MPECs ($KLRG1^{lo}CD127^{hi}$) in WT and SATB1-Tg mice; Mean proportions of splenic $D^bNP_{366}$- and $D^bPA_{224}$-specific CD8 T cells expressing (**e**) PD-1 and (**f**) CX3CR1. (**g-j**) The proportion (**g, i**) and mean fluorescence intensity (**h, j**) of Granzyme $A^+$ $D^bNP_{366}$- and $D^bPA_{224}$-specific $CD8^+$ T cells isolated from the BAL (**g, h**) and spleen (**i, j**). Data is representative of two independent experiments with N=5 female B6 mice and N=5 female SATB1-Tg mice. Data represented as mean ± SEM. * indicates $p$-value≤0.05, ** $p$≤0.01, *** $p$≤0.001 and **** $p$≤0.0001, unpaired student's $t$-test.

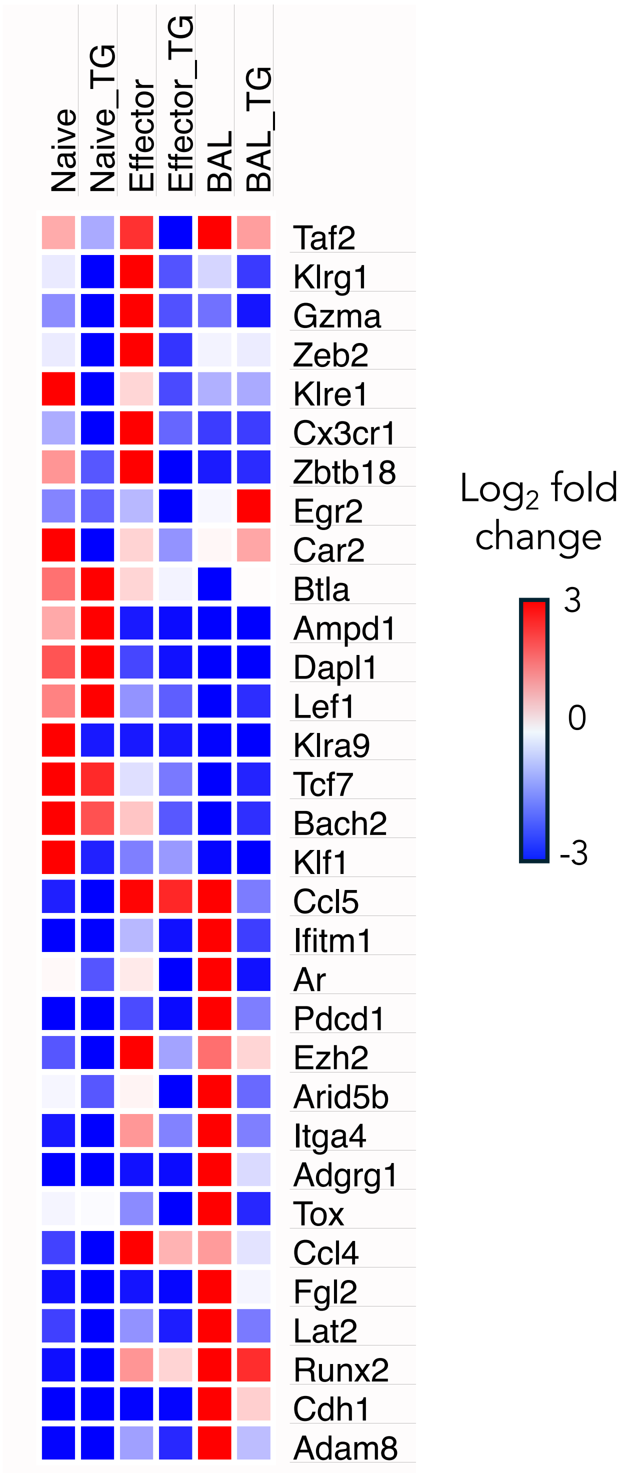


**Supplementary Figure 6.** Heatmap showing selected DEGs between WT and SATB1-Tg OT-I T cells. Naïve, and effector (BAL and Spleen) WT and SATB1-Tg OT-I CD8$^{+}$ T cells were isolated and RNA extracted for RNA-sequencing. DEGs were determined using DEGUST and the log2 fold change in transcript levels represented by heat map.
